# Effect of Substrate Stiffness on Human Intestinal Enteroids Infectivity by Enteroaggregative *Escherichia coli*

**DOI:** 10.1101/2021.06.15.448533

**Authors:** Ganesh Swaminathan, Nabiollah Kamyabi, Hannah E. Carter, Anubama Rajan, Umesh Karandikar, Zachary K. Criss, Noah F. Shroyer, Matthew J. Robertson, Cristian Coarfa, Chenlin Huang, Tate E. Shannon, Madeleine Tadros, Mary K. Estes, Anthony W. Maresso, K. Jane Grande-Allen

## Abstract

Human intestinal enteroids (HIE) models have contributed significantly to our understanding of diarrheal diseases and other intestinal infections, but their routine culture conditions fail to mimic the mechanical environment of the native intestinal wall. Because the mechanical characteristics of the intestine significantly alter how pathogens interact with the intestinal epithelium, we used different concentrations of polyethylene glycol (PEG) to generate soft (∼2 kPa), medium (∼10 kPa), and stiff (∼100 kPa) hydrogel biomaterial scaffolds. The height of HIEs cultured in monolayers atop these hydrogels was 18 µm whereas HIEs grown on rigid tissue culture surfaces (with stiffness in the GPa range) were 10 µm. Substrate stiffness also influenced the amount of enteroaggregative *E. coli* (EAEC strain 042) adhered to the HIEs. We quantified a striking difference in adherence pattern; on the medium and soft gels, the bacteria formed clusters of >100 and even >1000 on both duodenal and jejunal HIEs (such as would be found in biofilms), but did not on glass slides and stiff hydrogels. All hydrogel cultured HIEs showed significant enrichment for gene and signaling pathways related to epithelial differentiation, cell junctions and adhesions, extracellular matrix, mucins, and cell signaling compared to the HIEs cultured on rigid tissue culture surfaces. Collectively, these results indicate that the HIE monolayers cultured on the hydrogels are primed for a robust engagement with their mechanical environment, and that the soft hydrogels promote the formation of larger EAEC aggregates, likely through an indirect differential effect on mucus.

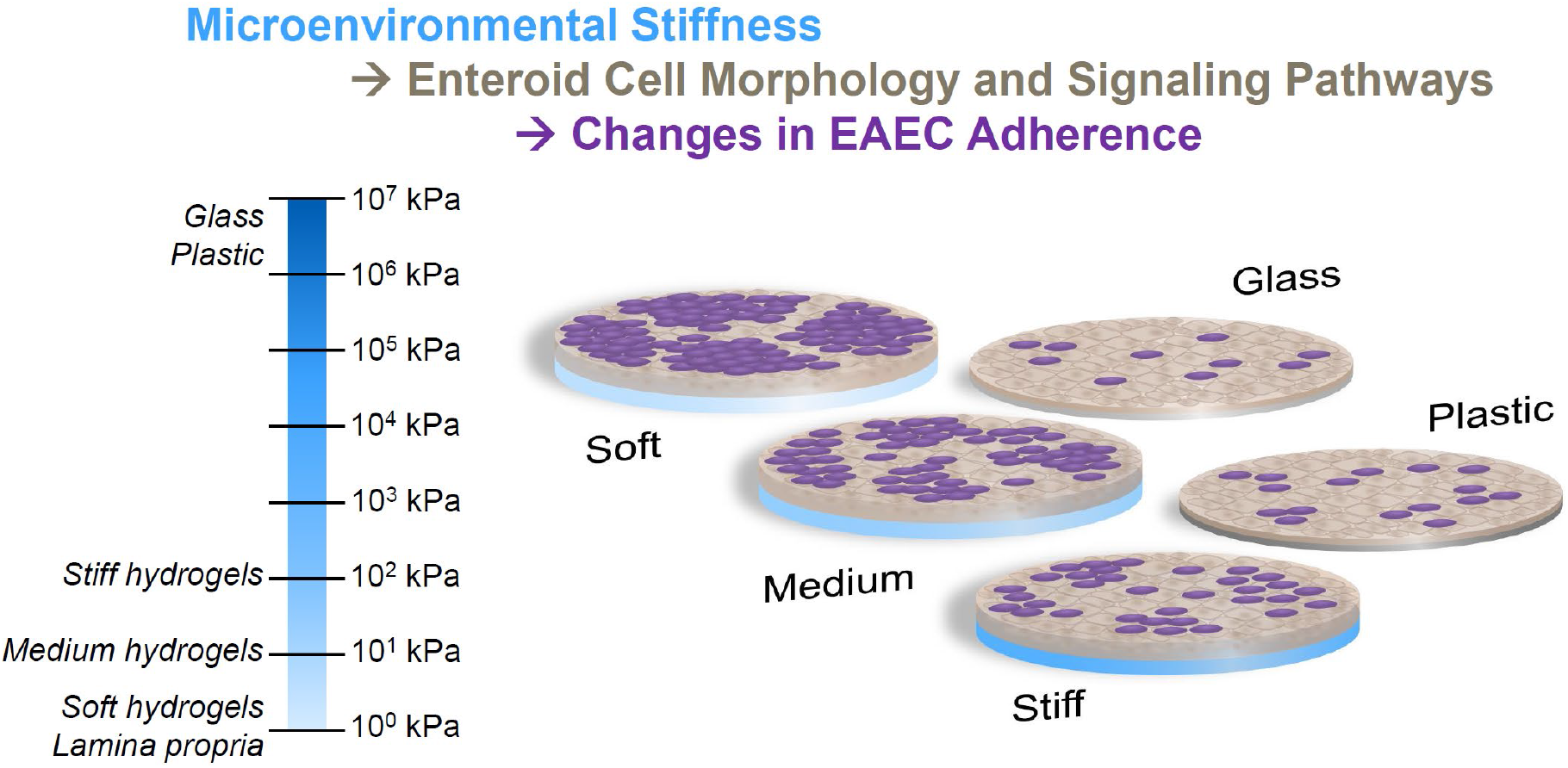
Graphical Abstract.

## 1. Introduction

Scientists have recently developed techniques to create human enteroid (enteric organoid) cultures as pre-clinical models to study the pathophysiology of enteric diseases as well as to develop drug therapies [1-3]. These 3D enteroid cultures, derived from intestinal stem cells in crypts isolated from human tissues (endoscopic biopsies and discarded surgical specimens), grow to include the entire crypt-villus axis, i.e., all epithelial cell types normally present. Enteroids naturally form in 3D with the apical surface of the epithelium located interiorly and the basal surface externally. This formation leads to difficulty in accessing the apical surface without invasive and time-consuming methods that are also not easily scalable for high throughput. These challenges have led investigators to continue to pursue further development of 2D models to support mechanistic studies with a variety of intestinal pathogens. One critical aspect to this model development is the consideration of the substrate mechanics, especially the stiffness.

Epithelial cells reside atop the mucosal layer (the *lamina propria*) of the villus, a loose connective tissue, and several layers of muscle (the submucosa and the muscularis mucosa). Despite the layers of smooth muscle and connective tissue, the layers directly in contact with the *lamina propria* and intestinal basement membrane are much softer. In contrast, the current culture substrates for 2D human intestinal enteroids (HIEs), Matrigel-coated plastic/glass, are extremely rigid. A more biomimetic culture platform for enteroids that closely mimics the soft nature of *lamina propria* can be provided by synthetic, customizable hydrogel scaffolds constructed using polyethylene glycol (PEG), which can be biofunctionalized by coating with proteins, peptides, or other bioactive molecules to mimic the ligand presentation of the basement membrane [4]. By controlling the degree of crosslinking, polymer molecular weight, or polymer volume fraction, one can control the hydrogel stiffness, which has been shown to influence the behavior of retinal epithelial cells cultured atop these gels [5]. PEG-based synthetic hydrogels containing cell-adhesive peptides were recently investigated as an alternative matrix for the culture and differentiation of intestinal organoids maintained in 3D culture [6, 7]. Notably, Gjorevski et al. [6] found that organoid expansion and differentiation were increased in stiffer and softer matrices, respectively.

This work was designed to investigate the impact of stiffness on HIE biological pathways and susceptibility to bacterial infection. Motivated by the need for improved access to the apical side of intestinal enteroids to study enteric infections, we recently reported a new approach to culture human intestinal epithelial enteroids as a monolayer on synthetic hydrogels [8]. Here, we examined the effect of hydrogel stiffness on HIEs cultured in 2D and then inoculated with Enteroaggregative *Escherichia coli* (EAEC). EAEC, named for its characteristic aggregative adherence pattern [9], is a global pathogen causing life-threatening diarrhea in developing nations [10], subclinical intestinal inflammation [11], long-term colonization in children associated with growth stunting, long term cognitive impairment, and malnutrition [12], and numerous cases of travelers’ diarrhea [13]. The aggregative phenotype of EAEC has been studied extensively in Hep2 cells [9], observed in experiments using pediatric intestinal explants [14] wherein EAEC formed 3D biofilm-like structures, and recently studied in HIEs [2]. The unique adherent and aggregative features of EAEC set the stage for the subsequent infection and disease symptoms with the prevailing model of pathogenesis beginning with adherence to the intestinal mucosa mediated by aggregative adherence fimbriae (AAF) [15, 16] and the involvement of AAF in the disruption of barrier integrity [17], the release of cytokines [18], and the induction of host inflammatory signaling [19] observed in EAEC infection.

The heterogeneity of clinical strains [20, 21] and lack of appropriate animal models [22] have complicated our understanding of EAEC pathogenic mechanisms, driving a need for high-throughput, biomimetic models of enteric infection. 3D HIE cultures have facilitated numerous discoveries about enteric pathogens [23], but we sought to develop a 2D model that would offer access to the apical surface as well as a customizable substrate stiffness that mimicked the mechanical behavior of the *lamina propria*. Thus, the goal of this work was to investigate the influence of substrate stiffness on human epithelial enteroids cultured in 2D monolayers and on their infection by EAEC.

## 2. Materials and Methods

### 2.1 Measurement of intestinal mucosal stiffness using micropipette aspiration

The small intestinal segments from young adult pigs (4-6 months; n = 3) were obtained from a local commercial abattoir (Animal Technologies, Tyler, TX). The *ex vivo* segments were rinsed gently to preserve mucosa, cut open longitudinally and then into smaller pieces of about 4-5 mm length, and stored in PBS at 4ºC. To measure the mucosal stiffness, micropipette aspiration was performed as previously described [24]. Briefly, glass micropipettes of diameter 60-70 μm were positioned perpendicularly to the intestinal tissue with direct access to aspirate the mucosal villi and crypt tissue into the lumen of the micropipette. Aspiration was performed by application of vacuum pressure through a syringe pump at a constant rate. The pressure was recorded using a pressure transducer operated by Labview software (National Instruments, Austin, TX). At each pressure level, an image of the micropipette with tissue aspirated into the lumen was captured to measure the aspiration length and pipette diameter (**Supplemental Fig. S1A**). The stiffness values were then calculated by fitting a finite-strain hyperelastic neo-Hookean constitutive model to the data as previously described [25].

### 2.2 Fabrication of hydrogels for seeding HIEs

Hydrogel precursor solutions were prepared by dissolving 20 kDa 8-arm PEG-norbornene (PEG-8N; JenKem Technology USA, Plano, TX) and 10 kDa 8-arm PEG-thiol (PEG-8T, JenKem Technology) in a visible light photoinitiation solution [26] consisting of HEPES buffered saline, pH 8.3 with 1% (v/v) triethanolamine (Acros Organics, Fair Lawn, NJ) and 10 µM eosin-Y disodium salt (Sigma Aldrich, St. Louis, MO). This precursor solution was prepared in three different formulations to achieve three different stiffnesses in the resulting hydrogel **(Table 1)**.

**Table 1.** Composition of the hydrogels with three different stiffnesses.

| Polymer | Soft | Medium | Stiff |
| --- | --- | --- | --- |
| PEG-8N | 3.5 mM | 4 mM | 12 mM |
| PEG-8T | 0.65 mM | 1.5 mM | 9 mM |

The hydrogels were polymerized using a ‘click’ reaction between norbornene and thiol groups with Eosin-Y as photoinitiator (**Fig. 1A**). To fabricate hydrogels, 4 µL of prepolymer solution was pipetted onto a Sigmacote® (Sigma Aldrich)-treated glass slide between 320 µm spacer molds, covered with a Sigmacote®-treated coverslip, then crosslinked using 150 kLux of full spectrum white light (UltraTow LED Floodlight, Northern Tool and Equipment, Burnsville, MN) for 3 min. The molds were then peeled off the plate leaving the hydrogel disks on top of the glass slides. Six (6)-mm diameter cloning cylinders were secured to the coverslips with silicone grease to create small column wells around each hydrogel disk. PBS was added to the columns to maintain hydrogel hydration (**Fig. 1B**). The gels were functionalized with ECM proteins in a 2-step process. First, PBS was aspirated and a 100 µL volume of 1:1 mixture of 2% Eosin-Y and thiolated poly(D-lysine) (PDL-SH) was added to each hydrogel column prior to crosslinking by 150 kLux of white light for 3 min (**Fig. 1 A**,**B**). The PDL-SH/Eosin-Y mixture was aspirated and the hydrogels were washed with PBS. A 100 µL volume of 1 mg/mL Matrigel in PBS was then added to each column, and the column well plates were incubated for seeding with HIEs the next day.

**Fig. 1.**
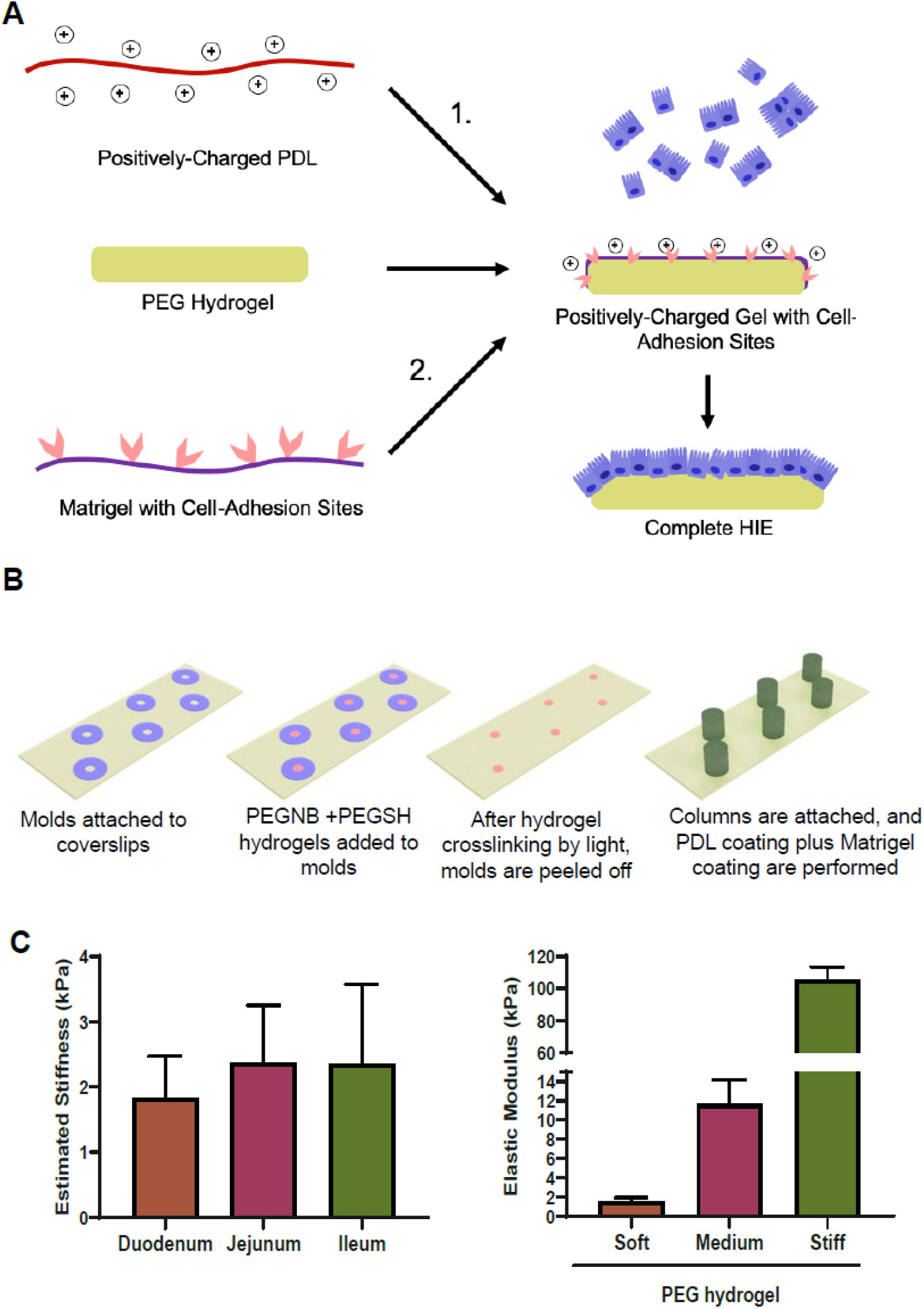
Hydrogel fabrication procedure for HIE culture. (A) Schematic of PEG hydrogel functionalization and HIE culture. (B) Schematic of various stages for PEG hydrogel fabrication on a glass slide using a PDMS mold and cloning columns. (C) Left, elasticity of intestinal wall segments measured by micropipette aspiration (n=3 per group). Right, elasticity of hydrogels (n=5 per group) measured using unconfined compression.

### 2.3 Mechanical testing of hydrogels

The three formulations of hydrogels were tested in uniaxial unconfined compression to determine their elastic moduli as described previously [27]. For this test, hydrogel disks were fabricated using the above precursor solutions at a set dimension of 6 mm diameter x 1 mm thickness using PDMS molds, crosslinked by white light, removed from the molds, and equilibrated in PBS overnight. Subsequently, the hydrogel disks were placed between the platens of a Bose ELF 3200 mechanical tester (Bose ELF, Eden Prairie, MN) and compressed until a strain of 40% was achieved. The reaction force was measured using a 1000 g load cell, while collecting force and displacement data. After converting the data to stress and strain by normalizing by the disk cross-sectional area and height, respectively, the compressive elastic modulus was calculated from a linear regression of the stress vs. strain plot between 5% and 15% strain. Three samples of the softest hydrogel formulations were also tested using micropipette aspiration as described in Section 2.1 (**Supplemental Fig. S1B**).

### 2.4 Isolation and culture of human intestinal epithelial enteroids (HIEs)

HIEs were established from human jejunal and duodenal epithelium and cultured as previously described [28]. Small intestinal biopsies were obtained from adults undergoing routine endoscopy or bariatric surgeries following a human subjects protocol approved by the Institutional Review Board at Baylor College of Medicine. All HIEs were derived either from human jejunum of three different individuals (J2, J3, J11) or duodenum of one individual (D109). Cultures of these established cell lines were grown in 3D forms in growth factor-reduced, phenol red-free Matrigel (Corning, Corning, NY) in complete medium with growth factors (CMGF+), which maintained the 3D HIEs in a predominantly undifferentiated, stem cell-rich state.

To obtain monolayer cultures of HIEs, undifferentiated 3D enteroids were released from the Matrigel with cold PBS containing 0.5 mM EDTA, pH 7.4, pelleted by centrifugation at 200 ×*g* for 5 min, and then incubated with trypsin for 4 min. After the trypsin was deactivated using medium with 10% serum (Fisher Scientific), cells were resuspended 4-5 times and strained (40 µm). The flow-through was collected, briefly pelleted (400 rcf for 5 min at room temperature), and then seeded atop the Matrigel-coated surfaces at a density of either 1×10^5^ cells/cm^2^ (96-well plates) or 3×10^5^ cells/cm^2^ (hydrogels) in CMGF+ medium supplemented with 10 µM Y-27632 (EMD Millipore) for 24 h to form monolayers. After the undifferentiated cells were adhered to the hydrogels or tissue culture plates, the cells were grown in differentiation medium, which contained CMGF+ medium without Wnt3A, SB202190, or nicotinamide and also contained 50% lower concentrations of Noggin and R-spondin [2, 29]. The HIE monolayers were cultured in differentiation medium for 5 days with medium changes every 48 h.

### 2.5 Assessment of morphology and differentiation with immunofluorescence and RT-qPCR

Immunofluorescent staining was used to determine the morphology of HIEs. Following culture for 5 days atop either the hydrogels of different stiffness or Matrigel-coated glass coverslips, the HIEs were fixed with 4% w/v paraformaldehyde (Fisher Scientific), permeabilized with 0.125% v/v Triton X-100 (Sigma Aldrich), and blocked with 5% v/v normal donkey serum (Abcam, Cambridge, MA). F-actin was visualized with Alexa Fluor-conjugated phalloidin (Invitrogen, Carlsbad, CA) and the nuclei were counterstained with DAPI (Invitrogen). Images were collected with confocal microscopy (Nikon A1-Rsi). Orthogonal slices of z-stack images were used to determine the height of the F-actin-stained cells using Fiji software. These cell heights were compared with measurements of epithelial cells in hematoxylin and eosin-stained sections of adult human jejunum and duodenum, obtained from our human subjects protocol as described in section 2.4.

To assess cell differentiation, additional samples of J2 HIEs cultured atop hydrogels were stained for beta-catenin, sucrose isomaltase (enterocytes), lysozyme (Paneth cells), and chromogranin A (enteroendocrine cells) as previously described in Wilson et al. [8] Phenotypic gene expression was assessed by reverse transcription quantitative polymerase chain reaction (RT-qPCR). Differentiated HIE monolayers from two individuals (J3 and D109) cultured atop hydrogels or tissue culture plastic lysed with Trizol (Thermo Fisher) and homogenized using a Tissue Lyser II (Qiagen). The cells on hydrogels were flash frozen prior to lysis. Total RNA was extracted using qScript XLT One-Step RT-qPCR ToughMix reagent with ROX reference dye (Quanta Biosciences). RT-qPCR was performed using TaqMan primer-probe mixes (Thermo Fisher) in a StepOnePlus system (Applied Biosystems) with glyceraldehyde-3-phosphate dehydrogenase (GAPDH) as the reference gene. Fold changes were calculated as 2^-ΔCT^.

### 2.6 Large-scale transcriptomics analysis

RNASeq analysis was performed to compare the gene expression of HIEs on PEG hydrogels with those on tissue culture plastic within 96-well plates. HIEs derived from the jejunum of 3 different individuals (J2, J3, J11) were seeded on 3 stiffnesses of PEG hydrogels (soft, medium, stiff) or on 96-well plates and cultured under differentiation conditions for 5 days as described above. For additional biological replication, RNA was collected from samples cultured on a range of dates, resulting in 36 samples total, including 3 replicates for each condition. After 5 days of differentiation culture, the hydrogels containing HIE monolayers were flash frozen, harvested in 1 mL of TRIzol (Thermo Fisher Scientific, Waltham, MA), and homogenized using a Tissue Lyser II (Qiagen, Germantown, MD). The cells on Matrigel-coated tissue culture plastic (96-well plates) were harvested directly in TRIzol. The harvested samples were transferred to new tubes and stored at -80ºC before being shipped for purification and mRNA-sequencing by Novogene (Sacramento, CA). The mRNA sequencing was performed using Illumina platforms for 150 bp paired-end reads.

The resulting fastq files were trimmed using the program wrapper for fastqc and cutadapt, TrimGalore! [30, 31]. Quality trimmed files were aligned against the human genome (GRCh38 p12) using the STAR aligner [32]. The human genome sequence and annotation file used during the alignment and subsequent read quantification were obtained from GENCODE [33]. Aligned reads were quantified using featureCounts at the gene level [34]. Protein coding gene counts for all samples were summarized into one matrix before performing differential gene analysis using the limma R-package with the voom transformation to correct for data heteroscedasticity [35]. For the linear models used in the differential gene analysis, gel stiffness was used as the explanatory variable and the patient from which the enteroids were derived was used as a blocking covariate. Pathway enrichment was determined using gene set enrichment analysis (GSEA) [36]. In the GSEA for each comparison considered, all detected genes were ranked according to their log2 fold change in expression. Heatmaps, volcano plots and principal component analysis (PCA) plots were created in the R-statistical computing environment.

### 2.7 Infecting HIEs with EAECs

HIE monolayers from four different patients (J2, J3, J11, and D109) were seeded on soft, medium, and stiff hydrogels, plastic 96 well plates (Ibidi, Fitchburg, WI), or glass chambered slides (Greiner Bio One, Monroe, NC) and differentiated for 5 or 6 days. Prototype EAEC strain 042 was grown overnight in tryptic soy broth (TSB) in a shaking incubator at 37°C. Prior to infection, 042 was subcultured for 3 h in fresh TSB at 37°C in a shaking incubator. Bacteria were then diluted in differentiation medium (3 µL per 100 µL fresh medium), which was added directly to the enteroids for infection. HIE monolayers were infected with 042 for 3 h at a multiplicity of infection (MOI) of 10 [37]. Neither DMEM nor the CMGF+ medium had a significant effect on the viability or growth curves of the bacteria (data not shown).

### 2.8 Assessment of infection using Giemsa staining/Colony Forming Unit (CFU) counting

#### Giemsa stain

Following infection, each well was washed 3x with room temperature PBS to remove unadhered bacteria. Slides were fixed and stained at room temperature using a Hema3 kit (Fisher Scientific). Representative images were taken at 100x magnification. Using these images, the number of bacteria and their pattern of adherence was counted [21].

#### CFU counting

Following infection, cells were washed gently 3x with room temperature PBS to remove unadhered bacteria. Cold PBS was added to each well to dissolve the Matrigel coating, and the cells (and bacteria adhered to them) were scraped off using a pipette tip. This solution was serially diluted in PBS, plated on luria broth plates, and grown overnight. Colonies were counted and the number of bacteria adhered to the cells in each well was calculated.

### 2.9 Statistical analyses

All statistical analyses were performed using one-way or two-way ANOVA with Tukey’s post-hoc tests unless stated otherwise. All values are reported as mean ± standard deviation with significance for p<0.05. Bacterial adherence assays (image analysis and CFU experiments) were analyzed using either ANOVA or Kruskal-Wallis’s test for non-parametric data with Dunn’s multiple comparison test. GraphPad Prism software was used for all analyses.

## 3. RESULTS

### 3.1 Micropipette aspiration demonstrates that the mucosal layer is soft

To provide a mechanical frame of reference for the development of hydrogel biomaterial scaffolds for the culture of HIEs, the stiffness of small intestinal segments from pigs was measured due to their anatomical and physiological similarity to human small intestine. To determine the stiffness of the mucosal layer, micropipette aspiration was performed. We found a trend of lower stiffness of the mucosal layer in the duodenum (1.84 ± 0.64 kPa) compared to the jejunum (2.38 ± 0.88 kPa) and ileum (2.35 ± 1.23 kPa), but this was not statistically significant **(Fig. 1C)**.

### 3.2 Synthetic hydrogels can be fabricated with a range of stiffness

To recapitulate the mucosal stiffness and to determine the effect of increase in stiffness on HIEs, we fabricated PEG-based hydrogels with three different stiffnesses by varying the weight fraction of the polymer constituents (**Table 1**). The elastic moduli measured from compressive mechanical testing were 1.58±0.36 kPa, 11.68±2.48 kPa, and 105.45±8 kPa for soft, medium, and stiff gels respectively **(Fig. 1C)**. The elastic modulus of the soft hydrogels measured using micropipette aspiration was 1.91 ± 0.41 kPa.

### 3.3 Confirmation of differentiation for HIEs cultured on hydrogels

We assessed whether the HIEs cultured atop the hydrogels were able to transition from a stem-like state into the multiple cell types found in differentiated intestinal epithelium. As shown in **Fig. 2A-C**, immunostaining for the epithelial marker beta-catenin demonstrated the integrity of the J2 HIE cell monolayer atop the hydrogels. The markers of differentiated enterocytes (sucrose isomaltase), Paneth cells (lysozyme), and enteroendocrine cells (chromogranin A) demonstrated that these cells were present in their expected low density (**Fig. 2D-L**). Duodenal (D109) HIEs grown on tissue culture plastic and on the hydrogels of different stiffnesses demonstrated no significant differences in gene expression of markers of enterocytes (*SI*), goblet cells (*MUC2*), enteroendocrine cells (*CHGA*), Paneth cells (*LYZ*), or intestinal epithelial stem cells (*HOPX, LRG5, CD44, KI67*) (**Fig. 2M**). Although the jejunal J3 line of HIEs showed the same trends in relative gene expression of these same markers (**Fig. 2N**), there were some significant differences in expression between the different substrate stiffness groups. These differences were generally minor, but there was unexpectedly high expression of the enterocyte markers *SI* and *ALPI* in the J3 HIE line cultured atop the soft hydrogels. Together, these results demonstrate that HIEs grown on these hydrogels of different stiffness retained their diverse range of cell phenotypes.

**Fig. 2.**
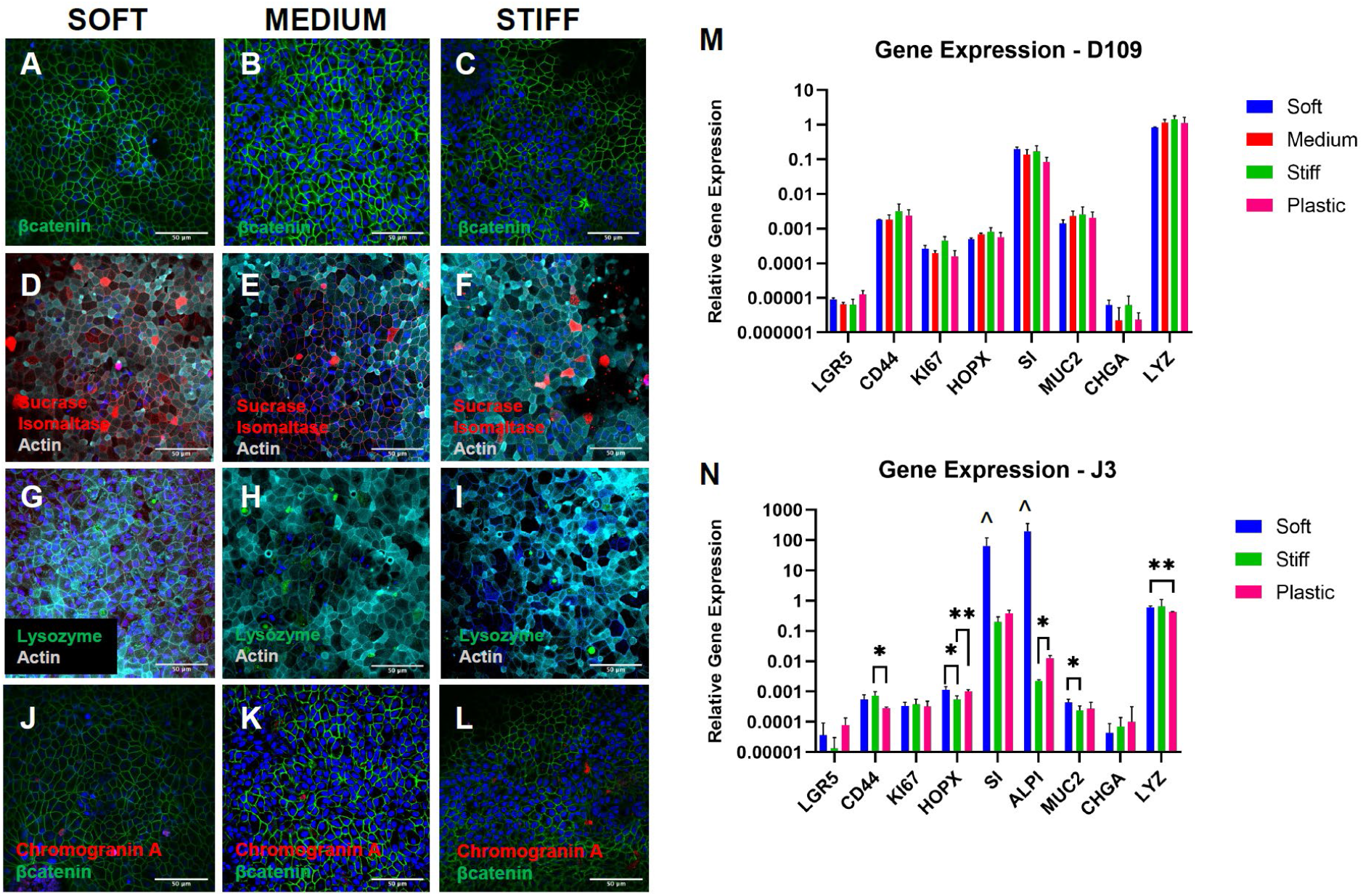
Confirmation of HIE phenotype. (A-L) Immunofluorescent staining of the differentiated J2 HIEs demonstrates the integrity of the cell monolayer (A-C) and phenotypic markers for three of the diverse cell types found in native intestinal epithelium: enterocytes (D-F), Paneth cells (G-I), and enteroendocrine cells (J-L). Blue in A-L: nuclei. Green in A-C and J-L: beta-catenin. Red in D-F: sucrose isomaltase. Teal in D-I: Actin. Green in G-I: Lysozyme. Red in J-L: Chromogranin A. Scale bar = 50 µm. (M-N) Analysis of the cellular composition of the HIEs by RT-qPCR reveals multi-lineage differentiation of duodenal and jejunal HIEs. Expression levels of each gene are show relative to GAPDH. There were no significant differences between duodenal D109 HIEs grown on hydrogels and cells grown on tissue culture plastic in 96 well plates (p>0.05, n=3 technical replicates). The J3 HIEs showed generally the same differentiation patterns, with some slight but significant differences between groups (p<0.05, n=6 technical replicates). Interestingly, the soft hydrogel/J3 cultures had unexpectedly strong expression of SI and ALPI, with a nearly significant trend of being greater than for the stiff hydrogel and plastic groups. The J3 medium hydrogel group had poor mRNA results for this data set. *p<0.05 between indicated groups, **p<0.01 between indicated groups, ^p=0.079 for SI and p=0.062 for ALPI vs. each of stiff and plastic.

### 3.4 Improved polarization of HIEs cultured on hydrogels

Previous studies have shown that stiffness of the culture substrate affects cell proliferation and differentiation in 3D intestinal organoids, however, it is unclear how stiffness will affect the morphology of HIEs grown in 2D. We tested the role of stiffness on cell morphology by comparing HIEs grown on coverglass to those grown on hydrogels of different stiffness, choosing to use the apical brush border and cell height as readouts. The polarization of differentiated HIEs was visualized by staining for F-actin, which is intensely expressed at the brush borders. HIEs on hydrogels with different stiffnesses were found to be present as flat monolayers that were appropriately polarized with brush borders, comparable to cells cultured on Matrigel-coated coverslips **(Fig. 3A)**. Interestingly, the height of HIEs cultured on hydrogels were approximately 80% taller (p<0.05) compared to HIEs cultured on glass coverslips (**Fig. 3B**), although still significantly shorter than epithelial cells in native human adult duodenal and jejunal tissues (**Fig. 3C-D**).

**Fig. 3.**
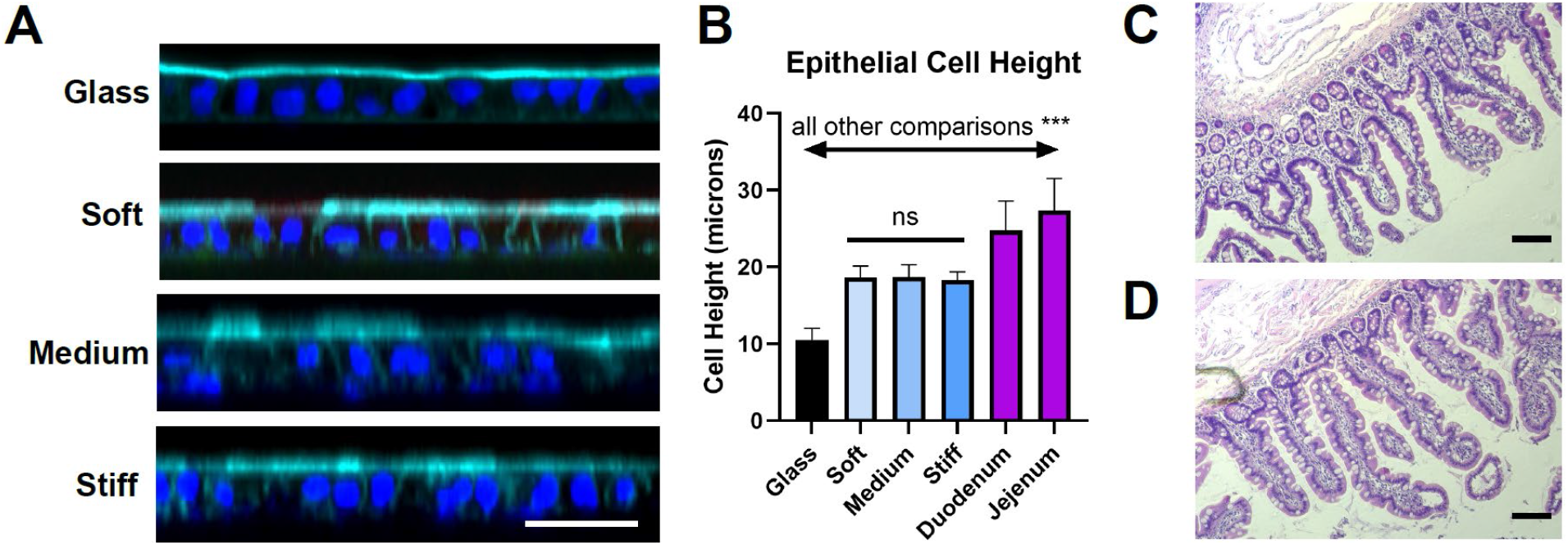
HIE cell height on hydrogels with various stiffness. (A) Confocal images depicting orthogonal view of HIE monolayers grown on various substrates. Te HIE cells appeared to be taller on the hydrogels compared with the glass coverslips. Teal = F-actin. Blue = nuclei. White scale bar = 50 µm. (B) Measurements of cell heights on glass coverslips, on hydrogels, and in tissues. Cell heights were not significantly different between the hydrogel groups, but all other comparisons were significantly different (***p<0.001). Each cell culture data set (17-32 total per group) was collected as 3-4 measurements from multiple distinct gels/coverslips. (C-D) Light microscopy images of (C) duodenum and (D) jejunum of human adult donor tissue (23 year old male). Tissue measurement shown in (B) were performed using ImageJ on 4 images per sample with 12-17 measurements per image. Black scale bars = 100 µm.

### 3.5 EAEC adherence was enhanced with reduced stiffness

To follow up on our previous report that EAEC adheres poorly to jejunal HIEs grown on glass [2], jejunal monolayers grown on various stiffness conditions and differentiated for 5-7 days were infected with prototype EAEC strain 042 following 3 h subculture of the bacteria in TSB. The bacteria did not appear to affect the integrity of the HIE monolayers, which was expected since EAEC is not an invasive pathogen. For EAEC, the relevant assessments are adherence and aggregation. We found that 042 adhered and aggregated in greater number on J2 monolayers grown with reduced stiffness (**Fig. 4A**). As the stiffness of plastic and glass are quite different from each other [38], we compared all hydrogels to both of these standard cell culture conditions.

**Fig. 4.**
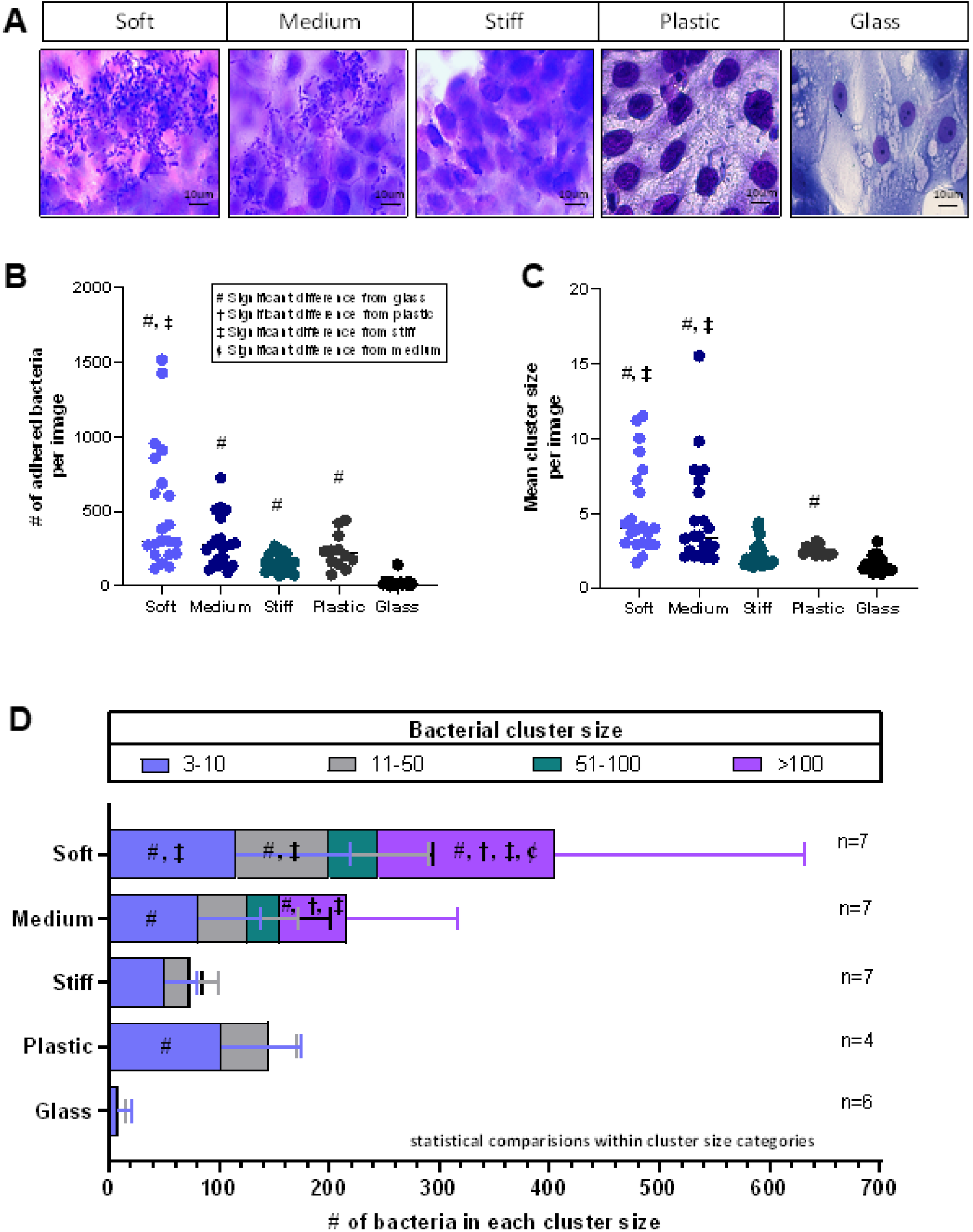
Adherence of Enteroaggregative E. coli to jejunal monolayers grown on hydrogels of various stiffnesses. (A) J2 HIE monolayers cultured with protype EAEC strain 042 for 3 hours, stained using a Hema3 kit and imaged at 100x magnification. Bacteria are small purple rods and nuclei are large purple ovals. (B-C) The total number of adhered bacteria (B) and number of bacteria in each cluster (C) were counted for each image. n represents the number of wells of each condition with 3 images taken per well (e.g., if n=6, 18 images were analyzed, 3 from each well). Each individual data point in (B) and (C) reflects the quantification of one image. (D) The mean number of bacteria in a given cluster size were compared across substrate categories. Data is from three separate experiments on different days. Cluster sizes of 1-2 bacteria are not shown in this graph. #, †, ‡, ¢ indicate significant difference from glass, plastic, stiff hydrogels, and medium hydrogels, respectively. Full tables of p-values for between-group comparisons are in the Supplementary Data.

We analyzed images of the J2 monolayers following infection and found an overall significant difference in number of adhered bacteria atop HIEs between the substrates of different stiffnesses (**Fig. 4B**, p<0.0001). The Dunn’s multiple comparison test (p-values provided in **Supplemental Table S1**) showed that all hydrogel groups as well as the plastic had significantly more adherent bacteria compared to the glass control (mean of 20.53 adhered bacteria/image): soft hydrogels (409, p<0.0001), medium hydrogels (168.2, p<0.0001), stiff hydrogels (57.20, p=0.0066), and plastic (231.3, p=0.0002). The soft vs. stiff hydrogel comparison was also significant (p=0.0016). Comparable data trends were also shown in studies of another jejunal line (J11) and a duodenal line (D109).

These findings were supported by quantification of colony forming units (CFUs) adhered to J11 monolayers under different stiffness conditions (**Fig. 5**). There was an overall significant difference in adhered CFUs between the different stiffnesses, p=0.0235, with a mean adhered CFU value of 3.00 × 10^5^ for glass, 5.93 × 10^5^ for stiff, 7.28 × 10^5^ for medium, and 7.65 × 10^5^ for soft. Dunn’s test for multiple comparisons revealed significant differences in mean adhered CFUs between the soft hydrogels (p=0.0177) and medium hydrogels (p=0.0402) compared to the glass control, but not between the stiff hydrogels and the glass control. As shown in **Supplemental Fig. S4**, CFU studies on the other cell lines gave comparable results with a non-significant trend of more adherent CFUs in the soft hydrogel group compared to the glass group for J2 (p=0.0703), no significant difference in CFUs for J3, and significantly more adherent CFUs in the soft group compared with the stiff hydrogel group for D109 (p=0.0443).

**Fig. 5.**
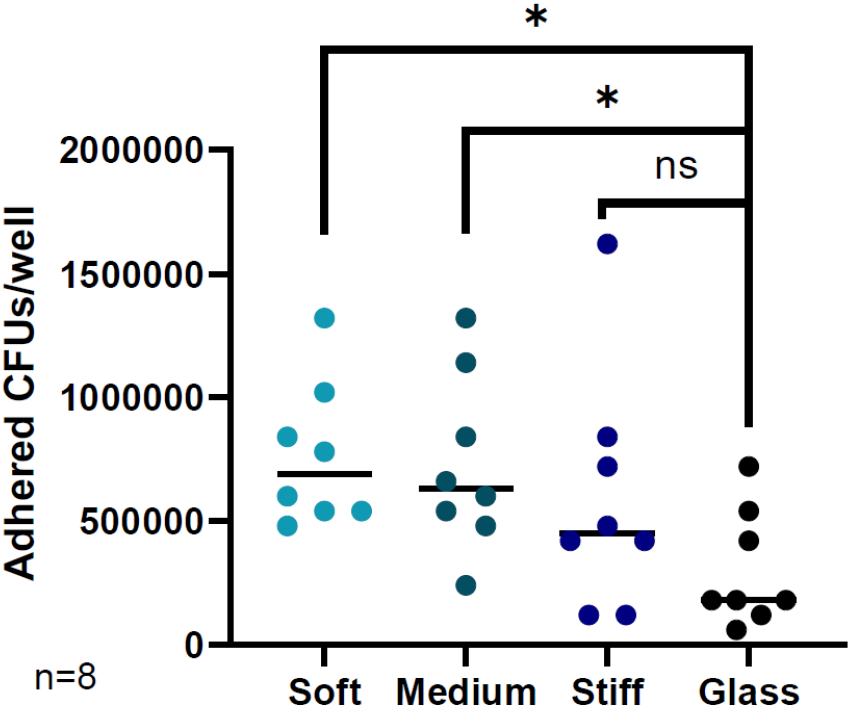
Adherence of Enteroaggregative E. coli to jejunal monolayers grown on hydrogels of various stiffnesses. J11 HIE monolayers were cultured with protype EAEC strain 042 for 3 hours, washed to remove unadhered bacteria, and scraped off the plate. The cell solution was then serially diluted, plated on LB plates, and counted for CFUs. Each data point represents the calculated CFU for one well. n=8 wells per condition. * denotes p≤0.05 vs. glass.

### 3.6 EAEC aggregation was enhanced with reduced stiffness

Additionally, we counted the size of each cluster of bacteria adhered to the HIEs [21] and found that EAEC aggregation (number of bacteria per cluster) increased with reduced stiffness. There was a significant difference in the mean cluster size (**Fig. 4C**) between various stiffnesses overall (p<0.0001). Dunn’s multiple comparison test revealed significant differences in mean cluster size on the soft hydrogels (p<0.0001), medium hydrogels (p<0.0001), and plastic (p=0.0242) compared to the glass control (all p-values provided in **Supplemental Table S2**). The mean cluster size was also greater on the soft hydrogels (p=0.0001) and medium hydrogels (p=0.0079) compared to the stiff hydrogels.

As shown in **Fig. 4D**, as stiffness decreased, both the total number of adhered bacteria and the number of bacteria in larger clusters increased. On J2 monolayers plated on traditional culture substrates, the largest cluster was 11 bacteria on glass and 48 bacteria on plastic (data not shown). In contrast, on gels of all three stiffnesses, EAEC formed clusters of 51-100; very few clusters of this size were observed on stiff gels whereas many were present on soft and medium gels. Only on soft and medium hydrogels were clusters greater than 100 bacteria observed. Statistically, the soft hydrogel cultures had the significantly greatest number of clusters of 1, 3-10, 11-50, and >100 bacteria (all p-values provided in **Supplemental Table S3**), whereas the glass cultures had the least. The medium hydrogel culture also had significantly more bacteria in clusters of >100 compared with the glass, plastic, and stiff hydrogel cultures. Comparable trends were also shown for the J11 and D109 lines. For the J11 line, HIEs atop both soft and stiff hydrogels had significantly more large clusters of adherent bacteria compared with HIEs cultured on glass (**Supplemental Fig. S2**). For the D109 line, HIEs atop the soft hydrogel had more adherent bacteria and larger bacterial clusters than the other groups, with some especially large colonies of >1000 bacteria (**Supplemental Fig. S3**).

### 3.7 RNASeq analysis shows substrate-driven differences

These differential results of varied EAEC adherence and aggregation to HIEs cultured atop substrates of distinct stiffness led us to hypothesize that biological signaling pathways, especially those relevant to mechanosensing and infection, would be altered by changes in the microenvironmental stiffness of the HIE cultures. Therefore, we performed a transcriptome analysis of HIEs cultured on soft, medium, and stiff hydrogels and 96-well plates. PCA showed pronounced differences in gene expression between the PEG hydrogels overall (green, purple, and blue symbols) and the 96-well plates (red dots) (**Fig. 6A**). Importantly and interestingly, we did not find differences in the gene expression of the HIEs when only comparing between the three gel stiffnesses. Volcano plots of the three hydrogels vs. 96-well comparisons demonstrated a roughly equivalent proportion of upregulated and downregulated genes (**Fig. 6B**). Of over 8000 profiled transcripts, for a log fold change (FC) of at least 2 with FDR < 0.05, ∼430-460 genes were upregulated and ∼350-410 genes were downregulated for each type of hydrogel compared to the 96-well plates. The majority (approximately 64%) of the significantly different genes were shared between the three hydrogels (**Fig. 6C**). All significantly different genes (for hydrogels vs. 96-well plates) can be found in **Supplemental Data Set S1**.

**Fig. 6.**
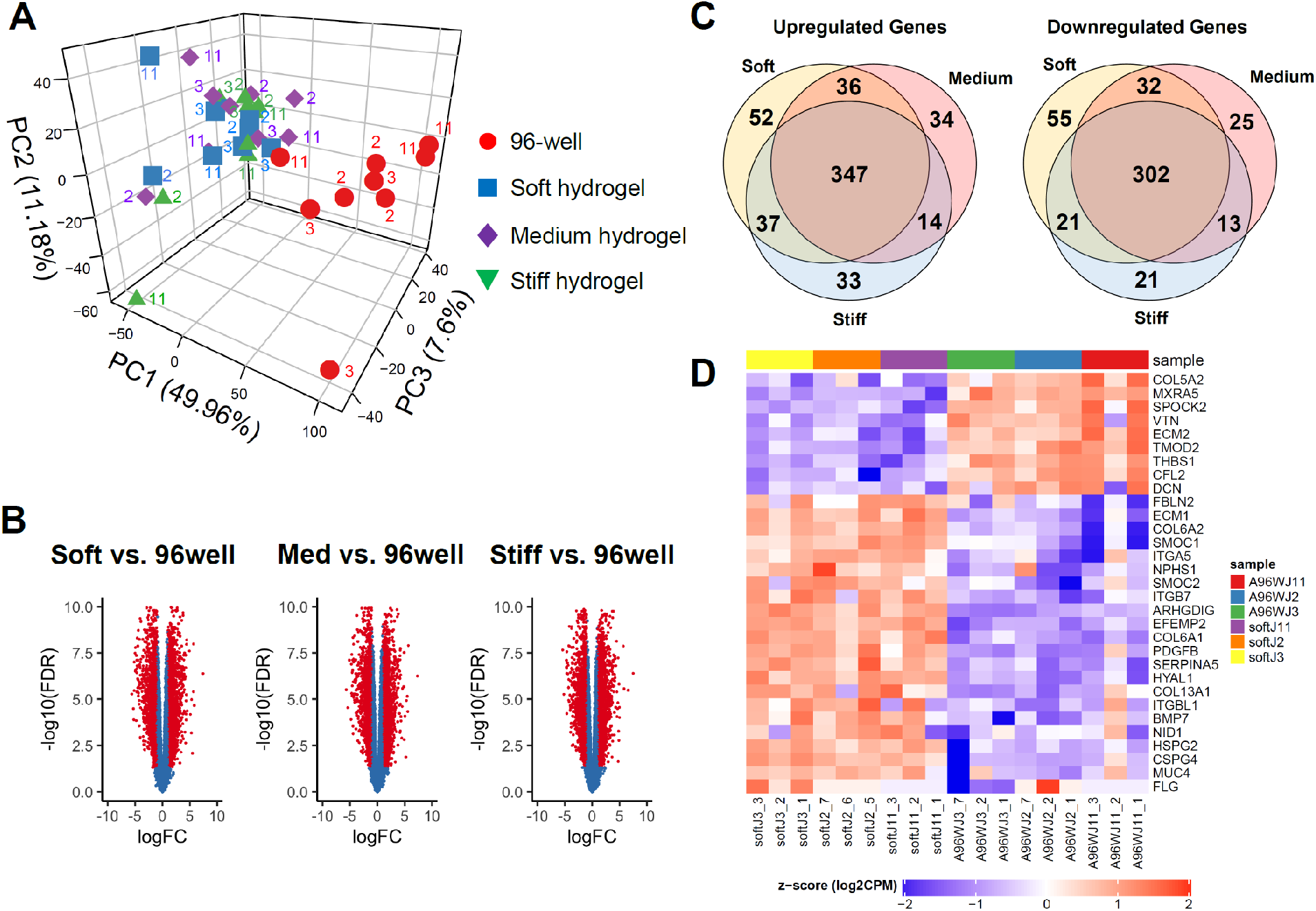
RNA sequencing analysis of human intestinal enteroid monolayers cultured on hydrogels or 96-well plates. (A) Principal component analysis of jejunal cultures demonstrating the variability observed in the samples (3 replicates of each substrate condition for 3 patient cell lines, J2, J3, and J11). (B) Volcano plot of differential gene expression for the three different gel stiffness compared to the 96-well controls. Red dots indicate genes that have an adjusted p-value of at least 0.05 with a log fold change (FC) of at least 2. (C) Venn diagrams present the number of genes that are differentially expressed (FC at least 2) within the cells cultured on soft, medium, and stiff gels compared to the 96-well cultures. Gene data are found in Supplemental Data Set 1. (D) Heat map for selected genes from biological pathways, known to be mechanosensitive, that were identified through Gene Set Enrichment Analysis. All genes shown have an FDR < 0.05 and log FC > 2. Left columns show gene expression in the soft hydrogels and right columns show the 96-well plates. Red = upregulated, blue = downregulated. The heat maps for the medium and stiff hydrogels vs. 96-well plates were very similar. This is a subset of the larger heat map provided in Supplemental Fig. S1. Data for the heatmaps is found in Supplemental Data Set 2.

Through a Gene Set Enrichment Analysis (GSEA), the dominant biological pathways correlated with the experimental groups were identified. Pathways with the greatest number of significant normalized enrichment scores (NES) with an FDR<0.05 were used to compile a heatmap of gene expression from several pathways that are known to be mechanosensitive [39], namely adhesion, extracellular matrix, external stimulus response, Rho GTPases, and cytoskeletal organization. The heatmap of all genes from these pathways demonstrating a log FC of at least 2.0 is shown in **Supplemental Fig. S5**, with a subset of these genes shown in **Fig. 6D**. These heatmaps show that culturing HIEs atop hydrogels with stiffness of approximately 2-100 kPa upregulates the majority of the genes in these pathways compared with HIEs on 96-well plastic plates. Of particular interest was upregulated gene expression of collagens (particularly types 6 and13), integrin subunits α5, β-like 1, and β7, mucins and proteoglycans, and basement membrane components such as nidogen, fibulins, and heparan sulfate proteoglycans in the hydrogel cultures. These heat maps and the significantly enriched pathway data can be found in **Supplemental Data Sets S2-S3**.

We examined several additional biological pathways to gain insight into the differential EAEC binding. With respect to cell differentiation, the cells cultured atop hydrogels showed strong enrichment of the Gene Ontology (GO) Epithelial Cell Differentiation and Canonical Wnt signaling pathways (**Supplemental Fig. S6A-B**). Therefore, epithelial differentiation and Wnt signaling genes with log FC>2 were compiled into a single heatmap (**Supplemental Fig. S6C**) with a subset heatmap in **Fig. 7A)**. HIEs cultured on gels had upregulation of several Wnt pathway genes including Frizzled Related Protein, NKD Inhibitor of WNT Signaling Pathway 2, and Wnt Family Members 4 and 7B. Similarly, epithelial differentiation genes Phosphodiesterase 2A, Sox8, and Prosaposin Like 1 were among the most upregulated in hydrogel cultures.

**Fig. 7.**
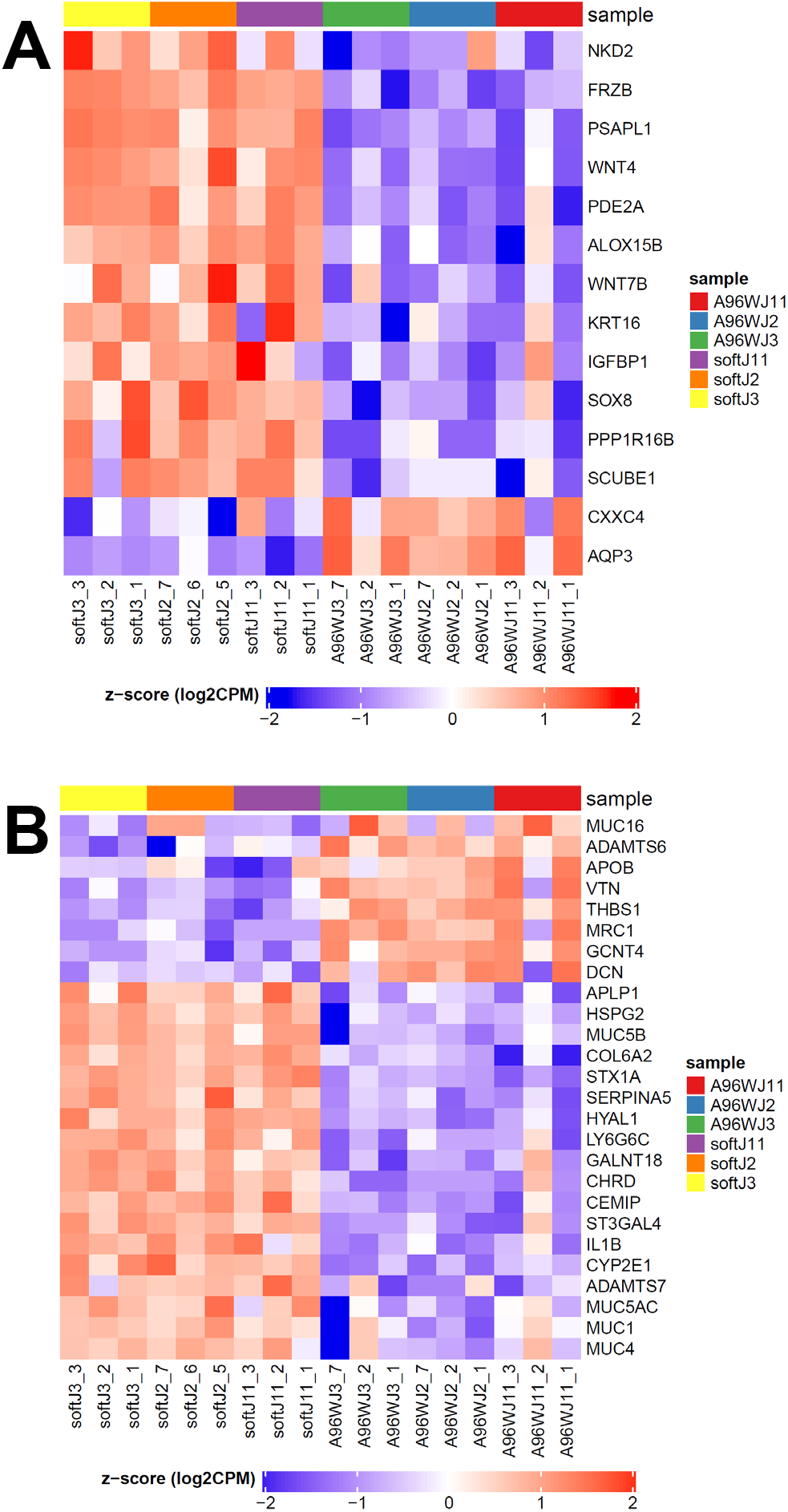
Heat maps of genes from biological pathways that were identified through Gene Set Enrichment Analysis. All genes shown have an FDR < 0.05 and log FC > 2.0. Left columns show gene expression in the soft hydrogels and right columns show the 96-well plates (3 replicates per patient for each substrate condition). Red = upregulated, blue = downregulated. The heat maps for the medium and stiff hydrogels vs. 96-well plates were very similar. These are subsets of the larger heat maps provided in Supplemental Figs. S2-S3. Data for the heatmaps is found in Supplemental Data Set 2. (A) Heat map of selected genes related to differentiated state. (B) Heat map of selected genes related to bacterial adhesion, especially via mucins and heparan sulfate proteoglycans.

Of relevance to the EAEC results, the GO Response to Bacterium pathway heatmap showed many differences between the cultures plated on the gels and 96-wells (**Supplemental Fig. S7A**). Since we previously reported that EAEC adherence to HIEs was dependent upon heparan sulfate proteoglycans (HSPGs) [40], we also prepared a heatmap of genes relevant to proteoglycans with log FC>2.0 as shown in **Supplemental Fig. S7B**, with a subset heatmap in **Fig. 7B**. These showed that the hydrogel cultures had the majority of significantly upregulated genes, including heparan sulfate proteoglycan 2 and mucins 1 and 4. Additional significantly different genes of interest (albeit with a |log FC|∼1) were the upregulated syndecans 1 and 3 and the downregulated heparanase, which degrades heparan sulfate glycosaminoglycan chains.

Examination of the entire GSEA revealed that pathways related to cell metabolism, digestion, and transcription were least enriched in gels relative to 96-well plates, and pathways related to ECM and the cell response to external stress, immunity and disease, cell cycle, Rho GTPases, adhesion, cell signaling, and NOTCH were most enriched (**Fig. 8**). When examining the GSEA for different stiffness gels only, transcriptional pathways were least enriched in the stiffer gel cultures (i.e., more correlated with medium and softer gels). In contrast, pathways related to cell metabolism, cell signaling, hemostasis, response to external stimulus, digestion, glycans, transport, and immunity were most enriched in the stiffer gel cultures (**Supplemental Fig. S8**).

**Fig. 8.**
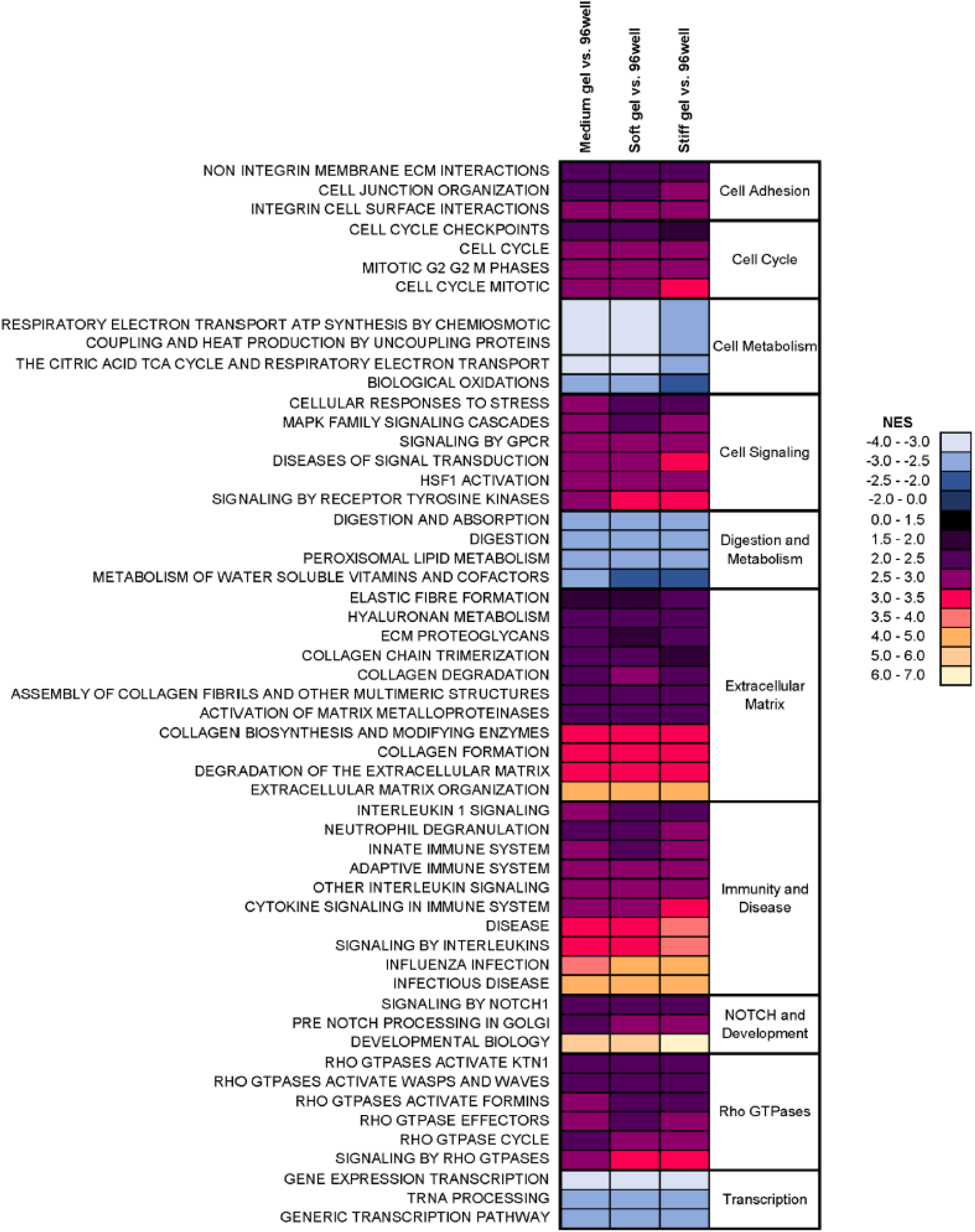
Normalized enrichment scores (NES) of selected Reactome pathways that were significantly different (FDR < 0.05) between HIEs cultured atop hydrogels and those cultured in 96-well plates. In each column, warmer colors indicate greater enrichment (positive correlation with gel) and cooler colors indicate least enrichment (positive correlation with 96-well plate). Data were averaged across the 3 patients.

## 4. DISCUSSION

The development of treatments for human diarrheal diseases has been complicated by the lack of reliable and relevant models for many enteric pathogens. Small animal models poorly mimic human disease pathophysiology, large animal models present challenges in terms of costs and scalability, and traditional *in vitro* models lack functional similarity with the human intestine [41]. Recently, the scientific community has applied 3D organotypic cultures, such as enteroids, for investigations of enteric infections since these cultures can recapitulate important aspects of intestinal cell phenotypes and functions [23]. Although most intestinal organoids and enteroids are cultured in 3D while surrounded by Matrigel, enteroid cultures can be converted to 2D monolayers on tissue culture plastic or glass (coated with a thin layer of Matrigel) to facilitate access to their apical surface. As both of these conditions differ mechanically from the native intestinal wall, they fail to mimic important mechanical aspects of host-pathogen interactions [42, 43]. Therefore, after measuring the stiffness of intestinal mucosa, we employed a hydrogel model with customizable stiffness to investigate the effect of substrate stiffness on the adherence and aggregation of enteroaggregative *E. coli* to primary human intestinal epithelial cells. After confirming that the HIEs cultured atop hydrogel scaffolds and those cultured on traditional tissue culture substrates were both successfully differentiated (as we recently reported [8]), we identified vast differences in HIE morphology and gene transcription between the two culture substrates. Further, we found that emulating physiological stiffness significantly enhances EAEC aggregation and adherence.

Even though three hydrogel stiffnesses were evaluated, the primary differences between the experimental HIE groups (uninfected) were between the traditional culture substrates and the hydrogels overall. This finding was observed for both the height of the HIE monolayer (taller for the hydrogels) as well as the large-scale transcriptional analysis. This difference in cell height cannot be attributed to microvilli alone, as those are typically one micron tall and the difference was several microns. Creff et al. recently showed a similar result with Caco-2 cells cultured atop flat 2D or villous-shaped 3D hydrogel being several microns taller than those cultured on glass [44]. Interestingly, the height difference in that study became slightly more pronounced after 21 days of culture, and the Caco-2 cell monolayers cultured on flat 2D hydrogels also spontaneously formed 3D villous-like structures. Although our HIE monolayers remained generally flat during the 5 days of differentiation culture, it would be intriguing to prolong this culture duration to assess whether a similar phenomenon occurs with the HIEs. As for the height difference, greater height of epithelial cells is an indicator of greater cell polarization and differentiation [45, 46], with positive correlations between cell height, robust junctional density, and cell integrity. The height of the differentiated HIEs cultured atop the hydrogels (approximately 18 µm) is closer to that of epithelial cells in tissue sections of the human small intestinal wall, which we measured as 25-27 µm. Correspondingly, there was significant enrichment for gene and signaling pathways related to epithelial differentiation and cell junctions in all hydrogel cultures. Together with the extensive enrichment of extracellular matrix components (including basement membrane) and cell-matrix adhesions (both integrin and non-integrin), these results indicate that the HIE monolayers cultured on the hydrogels are primed for a robust engagement with their mechanical environment. In contrast, the HIEs cultured on glass or tissue culture plastic (with stiffnesses in the GPa range [47]) were shorter and had more enrichment in signaling pathways related to transcription and cell metabolism. There was a pronounced stiffness-dependent effect on the adherence of EAEC to jejunal HIEs. We previously reported that EAEC 042 adheres poorly to jejunal HIE monolayers (cultured on glass), predominantly displaying a diffuse pattern of adherence [2]; this poor adherence is found with both differentiated and undifferentiated HIEs cultured on glass (data not shown). When J2 HIEs were grown on lower stiffness hydrogels for this work, the adherence pattern of 042 changed, forming larger 2D and 3D aggregates with increased adherence. We found these same striking results with a duodenal cell line, D109, and both stiff and soft hydrogels promoted this result for a different jejunal line, J11. However, when we investigated Transwell membranes (which are also less stiff than plastic and glass [48]), we did not find any difference in adherence of EAEC 042 to jejunal monolayers on Transwells compared to those on glass and plastic (**Supplemental Figs. S9-S10**).

Several possible explanations for increased EAEC adherence to HIEs on soft hydrogels were provided by the RNASeq data. The association between intestinal mucus and EAEC adherence has been known for decades [14], but the mechanism remains elusive. These jejunal HIEs on soft hydrogels had upregulation of the cell surface-associated mucins MUC1 and MUC4, and gel-forming mucins MUC5AC and MUC5B. EAEC has previously been linked with increased MUC1 expression [19] and found to interact with MUC1 in an AAF-dependent manner [19], suggesting a role for MUC1 as an intestinal ligand for EAEC. Additionally, we recently reported that EAEC’s major fimbrial subunit of aggregative adherence fimbria II (AafA) binds to heparan sulfate moieties [40]. This link is consistent with our finding of upregulation of heparan sulfate proteoglycan 2 (HSPG2) in the gel cultures as well as the genes for glycosaminoglycan biosynthesis and modification. We also noted upregulation of many genes for ECM components (as previously reported to [49] accompany EAEC infection), which provide a further source of binding to HSPGs [49]. We propose that heparan sulfate polysaccharides are key to these interactions between EAEC, intestinal mucus, and other cell-surface and extracellular components. Furthermore, we assert that the investigation of EAEC adherence to HIEs was transformed through the hydrogels’ mimicry of intestinal physiological stiffness. These findings should guide future studies to understand specific host-pathogen interactions at mucosal surfaces in the HIE model, especially given the lack of a feasible animal model for enteric pathogens.

Hydrogels can be prepared with a range of stiffnesses, and the GSEA revealed many compelling differences in enrichment of specific pathways between the soft, medium, and stiff gels. Although the comprehensive pathway data set showed significant enrichment of numerous pathways in the Gene Ontology, Reactome, KEGG, Hallmark, and other databases, we analyzed the Reactome pathways in more detail because they were concise yet representative of the whole GSEA. When comparing the enriched Reactome pathways from the gels-vs.-96 well plates comparison with the between-gels comparison, the pathways related to cell metabolism and digestion/metabolism were more enriched in the 96-well plate cultures, and also more enriched in the stiffer gels. This is a consistent pattern showing an effect of greater stiffness regardless of the substrate. In other instances, certain pathways (such as GPCR signaling, ECM organization, and innate immunity) were more enriched in the hydrogels overall compared to the 96-well plates, and then more enriched in the stiff gels compared to the medium and soft gels. Similarly, transcription pathways were least enriched in both the hydrogels overall as well as in the stiff gels in the between-gels comparison. These comparison patterns suggest that there may be a “sweet spot” of optimal pathway enrichment in a specific range of stiffness values (that includes the magnitude 100 kPa) for certain biological phenomena. A similar “sweet spot” of PEG hydrogel stiffness (190 Pa) was reported to promote intestinal stem cell proliferation via YAP signaling ([6]; in that model, the stem cells were maintained as 3D organoids encapsulated within the scaffold. Indeed, we found that three signaling pathways related to YAP (Gene Ontology hippo signaling, Reactome RUNX3 regulation of YAP1-mediated transcription, and Cardenonsi YAP conserved signature) were enriched in the stiff hydrogels relative to the medium or soft gels. Given the role of YAP in mechanosensation, regulation of canonical Wnt signaling, and regulation of secretory differentiation, it will be important in the future to investigate links between YAP, differentiation of specific intestinal cell subpopulations, mucus production, and EAEC infectivity. Other investigators have shown how synthetic biomaterials or intestinal stromal tissue equivalents fabricated with villous topography affect intestinal stem cell renewal, cell proliferation, and cell function [50-52]. In a similar vein, intestinal cells cultured on villous-patterned biomaterial scaffolds and exposed to shear stresses were shown to produce more mucins [53]. Our collaborative group is currently conducting a comprehensive analysis of dozens of transcriptomics analyses of HIEs from multiple intestinal segments across a range of culture substrates of different stiffnesses, including tissue culture plastic, soft Matrigel, hydrogels, and Transwell inserts [48].

Lastly, micropipette aspiration was used to quantify the stiffness of the *lamina propria*, which at 1-3 kPa is profoundly less stiff than the underlying connective tissue (approximately 25 MPa [54]). While the stiffness measurements of the soft hydrogel are comparable between compression testing and micropipette aspiration (**Supplementary Fig. S5**), the working limit of our micropipette aspiration system is less than 5 kPa. The low stiffness of the mucosa provides context for the mechanical milieu of the intestinal epithelium: cells growing on the villi are adhered to a soft, easily deformable tissue. In contrast, the cells located in the crypts are also adhered to this soft tissue but may also be subject to an additional source of mechanically-based signaling due to their closer proximity to the much stiffer submucosa. In the future, it will be compelling to investigate how the heterogeneous intestinal wall provides a range of mechanical stimulation to the intestinal epithelial cells.

Although these results demonstrating the effect of substrate stiffness on the binding of EAEC to HIEs are very compelling, they must be considered in context. These experiments were performed with HIEs only, and therefore the additional cell types that make up the intestinal wall community were not present, such as intestinal fibroblasts, microvascular cells, and commensal microbiota. Interactions between these cells may be affected by substrate stiffness, and those interactions may correspondingly impact adherence and aggregation by EAEC. Additionally, the hydrogels themselves presented challenges in histological processing, such as for dehydration, paraffin embedding, and sectioning for standard histology and for the drying steps needed for electron microscopy (for demonstration of microvilli). Mucus and mucins in particular are notoriously challenging to image, and some of the fixatives needed for mucus staining are rather corrosive and could not be used with the hydrogels. We were also concerned that other mucus stains might be retained by the hydrogels. We are continuing to optimize our methods for the staining of cells cultured on these hydrogels. Furthermore, it was not possible to measure trans-epithelial electrical resistance (TEER) of the HIEs in these hydrogel cultures, although we have previously reported successful measurement of TEER for HIEs cultured atop Transwells [55]. Although EAEC is not an invasive pathogen, future studies could readily be envisioned in which invasive bacteria are expected to disrupt the layer of epithelial cells grown atop the hydrogel. In such cases, it will be essential to develop or modify a means of measuring functional integrity of the epithelial barrier. In conclusion, we used a hydrogel model that mimics the stiffness of intestinal mucosa to demonstrate differences between HIEs cultured on hydrogels vs. tissue culture plates. Moreover, there was a stiffness-dependent effect on the adherence and aggregation of enteroaggregative *E. coli* to the HIEs, with the softer, more physiological hydrogels promoting significantly greater bacterial adhesion. These results are likely due to upregulation of mucins and other extracellular matrix components that engage with bacterial adhesive filaments. Further investigation of these phenomena is necessary to elucidate the role of intestinal mechanobiology in enteric disease.

## Supporting information

Supplemental Dat Set 1

Supplemental Data Set 2

Supplemental Data Set 3

## Acknowledgements

We thank Drs. Sarah Blutt and Sasirekha Ramani for helpful discussions about RNASeq data analysis and interpretation, Xiaomin Yu for technical assistance, and Luis Victor for assistance with the graphical abstract. This research was supported in part by NIH Grant U19 AI116497, the Texas Medical Center Digestive Diseases Center supported by NIH P30 DK56338, and the Advanced Technology Core Laboratories (Baylor College of Medicine), specifically the Integrated Microscopy Core with funding from CPRIT (RP150578, RP170719), The Dan L. Duncan Comprehensive Cancer Center, and Baylor College of Medicine Office of Research.

## Appendix A. Supplementary Data

Supplemental Figures S1-S10, Tables S1-S7, and Data Sets S1-S3 are available online.

## References

[1] J.R. Spence, C.N. Mayhew, S.A. Rankin, M.F. Kuhar, J.E. Vallance, K. Tolle, E.E. Hoskins, V.V. Kalinichenko, S.I. Wells, A.M. Zorn, N.F. Shroyer, J.M. Wells, Directed differentiation of human pluripotent stem cells into intestinal tissue in vitro, Nature 470 (2011) 105–9.

[2] A. Rajan, L. Vela, X.L. Zeng, X. Yu, N. Shroyer, S.E. Blutt, N.M. Poole, L.G. Carlin, J.P. Nataro, M.K. Estes, P.C. Okhuysen, A.W. Maresso, Novel segment- and host-specific patterns of enteroaggregative Escherichia coli adherence to human intestinal enteroids, mBio 9 (2018) e02419–17.

[3] J. Foulke-Abel, J. In, O. Kovbasnjuk, N.C. Zachos, K. Ettayebi, S.E. Blutt, J.M. Hyser, X.L. Zeng, S.E. Crawford, J.R. Broughman, M.K. Estes, M. Donowitz, Human enteroids as an ex-vivo model of host-pathogen interactions in the gastrointestinal tract, Exp Biol Med 239 (2014) 1124–34.

[4] C.R. Marti-Figueroa, R.S. Ashton, The case for applying tissue engineering methodologies to instruct human organoid morphogenesis, Acta Biomater 54 (2017) 35–44.

[5] C. White, T. DiStefano, R. Olabisi, The influence of substrate modulus on retinal pigment epithelial cells, J Biomed Mater Res A 105 (2017) 1260–1266.

[6] N. Gjorevski, N. Sachs, A. Manfrin, S. Giger, M.E. Bragina, P. Ordonez-Moran, H. Clevers, M.P. Lutolf, Designer matrices for intestinal stem cell and organoid culture, Nature 539 (2016) 560–564.

[7] V. Hernandez-Gordillo, T. Kassis, A. Lampejo, G. Choi, M.E. Gamboa, J.S. Gnecco, A. Brown, D.T. Breault, R. Carrier, L.G. Griffith, Fully synthetic matrices for in vitro culture of primary human intestinal enteroids and endometrial organoids, Biomaterials 254 (2020) 120125.

[8] R.L. Wilson, G. Swaminathan, K. Ettayebi, C. Bomidi, X.L. Zeng, S.E. Blutt, M.K. Estes, K.J. Grande-Allen, Protein-functionalized poly(ethylene glycol) hydrogels as scaffolds for monolayer organoid culture, Tissue Eng Part C Methods 27 (2021) 12–23.

[9] J.P. Nataro, J.B. Kaper, R. Robins-Browne, V. Prado, P. Vial, M.M. Levine, Patterns of adherence of diarrheagenic Escherichia coli to hep-2 cells, Pediatr Infect Dis J 6 (1987) 829–31.

[10] F. Qadri, A.M. Svennerholm, A.S. Faruque, R.B. Sack, Enterotoxigenic Escherichia coli in developing countries: Epidemiology, microbiology, clinical features, treatment, and prevention, Clin Microbiol Rev 18 (2005) 465–83.

[11] T.S. Steiner, A.A. Lima, J.P. Nataro, R.L. Guerrant, Enteroaggregative Escherichia coli produce intestinal inflammation and growth impairment and cause interleukin-8 release from intestinal epithelial cells, J Infect Dis 177 (1998) 88–96.

[12] R.L. Guerrant, M.D. DeBoer, S.R. Moore, R.J. Scharf, A.A. Lima, The impoverished gut--a triple burden of diarrhoea, stunting and chronic disease, Nat Rev Gastroenterol Hepatol 10 (2013) 220–9.

[13] M. Glandt, J.A. Adachi, J.J. Mathewson, Z.D. Jiang, D. DiCesare, D. Ashley, C.D. Ericsson, H.L. DuPont, Enteroaggregative Escherichia coli as a cause of traveler’s diarrhea: Clinical response to ciprofloxacin, Clin Infect Dis 29 (1999) 335–8.

[14] S. Hicks, D.C. Candy, A.D. Phillips, Adhesion of enteroaggregative Escherichia coli to pediatric intestinal mucosa in vitro, Infect Immun 64 (1996) 4751–60.

[15] J.R. Czeczulin, S. Balepur, S. Hicks, A. Phillips, R. Hall, M.H. Kothary, F. Navarro-Garcia, J.P. Nataro, Aggregative adherence fimbria ii, a second fimbrial antigen mediating aggregative adherence in enteroaggregative Escherichia coli, Infect Immun 65 (1997) 4135–45.

[16] A. Rajan, H. Shi, B. Xue, Class i and ii histone deacetylase inhibitors differentially regulate thermogenic gene expression in brown adipocytes, Sci Rep 8 (2018) 13072.

[17] M.C. Strauman, J.M. Harper, S.M. Harrington, E.J. Boll, J.P. Nataro, Enteroaggregative Escherichia coli disrupts epithelial cell tight junctions, Infect Immun 78 (2010) 4958–64.

[18] S.M. Harrington, M.C. Strauman, C.M. Abe, J.P. Nataro, Aggregative adherence fimbriae contribute to the inflammatory response of epithelial cells infected with enteroaggregative Escherichia coli, Cell Microbiol 7 (2005) 1565–78.

[19] E.J. Boll, J. Ayala-Lujan, R.L. Szabady, C. Louissaint, R.Z. Smith, K.A. Krogfelt, J.P. Nataro, F. Ruiz-Perez, B.A. McCormick, Enteroaggregative Escherichia coli adherence fimbriae drive inflammatory cell recruitment via interactions with epithelial muc1, mBio 8 (2017) e00717–17.

[20] N. Boisen, M.T. Osterlund, K.G. Joensen, A.E. Santiago, I. Mandomando, A. Cravioto, M.A. Chattaway, L.A. Gonyar, S. Overballe-Petersen, O.C. Stine, D.A. Rasko, F. Scheutz, J.P. Nataro, Redefining enteroaggregative Escherichia coli (EAEC): Genomic characterization of epidemiological EAEC strains, PLoS Negl Trop Dis 14 (2020) e0008613.

[21] A. Olvera, H. Carter, A. Rajan, L.G. Carlin, X. Yu, X.L. Zeng, S. Shelburne, M. Bhatti, S.E. Blutt, N.F. Shroyer, R. Jenq, M.K. Estes, A. Maresso, P.C. Okhuysen, Enteropathogenic Escherichia coli infection in cancer and immunosuppressed patients, Clin Infect Dis (2020) in press.

[22] C.W. Philipson, J. Bassaganya-Riera, R. Hontecillas, Animal models of enteroaggregative Escherichia coli infection, Gut Microbes 4 (2013) 281–91.

[23] W.Y. Zou, S.E. Blutt, S.E. Crawford, K. Ettayebi, X.L. Zeng, K. Saxena, S. Ramani, U.C. Karandikar, N.C. Zachos, M.K. Estes, Human intestinal enteroids: New models to study gastrointestinal virus infections, Methods Mol Biol 1576 (2019) 229–247.

[24] M.M. Fahrenholtz, S. Wen, K.J. Grande-Allen, Development of a heart valve model surface for optimization of surface modifications, Acta Biomater 26 (2015) 64–71.

[25] R. Zhao, K.L. Sider, C.A. Simmons, Measurement of layer-specific mechanical properties in multilayered biomaterials by micropipette aspiration, Acta Biomater 7 (2011) 1220–7.

[26] C.C. Lin, C.S. Ki, H. Shih, Thiol-norbornene photo-click hydrogels for tissue engineering applications, J Appl Polym Sci 132 (2015) 41563.

[27] D.S. Puperi, R.W. O’Connell, Z.E. Punske, Y. Wu, J.L. West, K.J. Grande-Allen, Hyaluronan hydrogels for a biomimetic spongiosa layer of tissue engineered heart valve scaffolds, Biomacromolecules 17 (2016) 1766–75.

[28] K. Saxena, S.E. Blutt, K. Ettayebi, X.L. Zeng, J.R. Broughman, S.E. Crawford, U.C. Karandikar, N.P. Sastri, M.E. Conner, A.R. Opekun, D.Y. Graham, W. Qureshi, V. Sherman, J. Foulke-Abel, J. In, O. Kovbasnjuk, N.C. Zachos, M. Donowitz, M.K. Estes, Human intestinal enteroids: A new model to study human rotavirus infection, host restriction, and pathophysiology, J Virol 90 (2016) 43–56.

[29] K. Ettayebi, S.E. Crawford, K. Murakami, J.R. Broughman, U. Karandikar, V.R. Tenge, F.H. Neill, S.E. Blutt, X.L. Zeng, L. Qu, B. Kou, A.R. Opekun, D. Burrin, D.Y. Graham, S. Ramani, R.L. Atmar, M.K. Estes, Replication of human noroviruses in stem cell-derived human enteroids, Science 353 (2016) 1387–1393.

[30] M. Martin, Cutadapt removes adapter sequences from high-throughput sequencing reads, EMBnet.journal 17 (2011) 3.

[31] A. S., Fastqc: A quality control tool for high throughput sequence data., 2010. http://www.bioinformatics.babraham.ac.uk/projects/fastqc/.

[32] A. Dobin, C.A. Davis, F. Schlesinger, J. Drenkow, C. Zaleski, S. Jha, P. Batut, M. Chaisson, T.R. Gingeras, Star: Ultrafast universal rna-seq aligner, Bioinformatics 29 (2013) 15–21.

[33] A. Frankish, M. Diekhans, A.M. Ferreira, R. Johnson, I. Jungreis, J. Loveland, J.M. Mudge, C. Sisu, J. Wright, J. Armstrong, I. Barnes, A. Berry, A. Bignell, S. Carbonell Sala, J. Chrast, F. Cunningham, T. Di Domenico, S. Donaldson, I.T. Fiddes, C. Garcia Giron, J.M. Gonzalez, T. Grego, M. Hardy, T. Hourlier, T. Hunt, O.G. Izuogu, J. Lagarde, F.J. Martin, L. Martinez, S. Mohanan, P. Muir, F.C.P. Navarro, A. Parker, B. Pei, F. Pozo, M. Ruffier, B.M. Schmitt, E. Stapleton, M.M. Suner, I. Sycheva, B. Uszczynska-Ratajczak, J. Xu, A. Yates, D. Zerbino, Y. Zhang, B. Aken, J.S. Choudhary, M. Gerstein, R. Guigo, T.J.P. Hubbard, M. Kellis, B. Paten, A. Reymond, M.L. Tress, P. Flicek, Gencode reference annotation for the human and mouse genomes, Nucleic Acids Res 47 (2019) D766–D773.

[34] Y. Liao, G.K. Smyth, W. Shi, Featurecounts: An efficient general purpose program for assigning sequence reads to genomic features, Bioinformatics 30 (2014) 923–30.

[35] C.W. Law, Y. Chen, W. Shi, G.K. Smyth, Voom: Precision weights unlock linear model analysis tools for rna-seq read counts, Genome Biol 15 (2014) R29.

[36] A. Subramanian, P. Tamayo, V.K. Mootha, S. Mukherjee, B.L. Ebert, M.A. Gillette, A. Paulovich, S.L. Pomeroy, T.R. Golub, E.S. Lander, J.P. Mesirov, Gene set enrichment analysis: A knowledge-based approach for interpreting genome-wide expression profiles, Proc Natl Acad Sci U S A 102 (2005) 15545–50.

[37] N.M. Poole, A. Rajan, A.W. Maresso, Human intestinal enteroids for the study of bacterial adherence, invasion, and translocation, Curr Protoc Microbiol 50 (2018) e55.

[38] S. Acevedo-Acevedo, W.C. Crone, Substrate stiffness effect and chromosome missegregation in hIPS cells, J Negat Results Biomed 14 (2015) 22.

[39] K. Ohashi, S. Fujiwara, K. Mizuno, Roles of the cytoskeleton, cell adhesion and rho signalling in mechanosensing and mechanotransduction, J Biochem 161 (2017) 245–254.

[40] A. Rajan, M.J. Robertson, H.E. Carter, N.M. Poole, J.R. Clark, S.I. Green, Z.K. Criss, B. Zhao, U. Karandikar, Y. Xing, M. Margalef-Catala, N. Jain, R.L. Wilson, F. Bai, J.M. Hyser, J. Petrosino, N.F. Shroyer, S.E. Blutt, C. Coarfa, X. Song, B.V. Prasad, M.R. Amieva, J. Grande-Allen, M.K. Estes, P.C. Okhuysen, A.W. Maresso, Enteroaggregative E. coli adherence to human heparan sulfate proteoglycans drives segment and host specific responses to infection, PLoS Pathog 16 (2020) e1008851.

[41] S.E. Blutt, J.R. Broughman, W. Zou, X.L. Zeng, U.C. Karandikar, J. In, N.C. Zachos, O. Kovbasnjuk, M. Donowitz, M.K. Estes, Gastrointestinal microphysiological systems, Exp Biol Med 242 (2017) 1633–1642.

[42] C.P. Gayer, M.D. Basson, The effects of mechanical forces on intestinal physiology and pathology, Cell Signal 21 (2009) 1237–44.

[43] R.G. Lentle, P.W. Janssen, Physical characteristics of digesta and their influence on flow and mixing in the mammalian intestine: A review, J Comp Physiol B 178 (2008) 673–90.

[44] J. Creff, R. Courson, T. Mangeat, J. Foncy, S. Souleille, C. Thibault, A. Besson, L. Malaquin, Fabrication of 3D scaffolds reproducing intestinal epithelium topography by high-resolution 3D stereolithography, Biomaterials 221 (2019) 119404.

[45] A. Vinaiphat, K. Charngkaew, V. Thongboonkerd, More complete polarization of renal tubular epithelial cells by artificial urine, Cell Death Discov 4 (2018) 47.

[46] G. Altay, E. Larranaga, S. Tosi, F.M. Barriga, E. Batlle, V. Fernandez-Majada, E. Martinez, Self-organized intestinal epithelial monolayers in crypt and villus-like domains show effective barrier function, Sci Rep 9 (2019) 10140.

[47] A. Skardal, D. Mack, A. Atala, S. Soker, Substrate elasticity controls cell proliferation, surface marker expression and motile phenotype in amniotic fluid-derived stem cells, J Mech Behav Biomed Mater 17 (2013) 307–16.

[48] M.J. Mondrinos, Y.S. Yi, N.K. Wu, X. Ding, D. Huh, Native extracellular matrix-derived semipermeable, optically transparent, and inexpensive membrane inserts for microfluidic cell culture, Lab Chip 17 (2017) 3146–3158.

[49] M.J. Farfan, K.G. Inman, J.P. Nataro, The major pilin subunit of the AAF/II fimbriae from enteroaggregative Escherichia coli mediates binding to extracellular matrix proteins, Infect Immun 76 (2008) 4378–84.

[50] C.M. Costello, J. Hongpeng, S. Shaffiey, J. Yu, N.K. Jain, D. Hackam, J.C. March, Synthetic small intestinal scaffolds for improved studies of intestinal differentiation, Biotechnol Bioeng 111 (2014) 1222–32.

[51] Y. Wang, D.B. Gunasekara, M.I. Reed, M. DiSalvo, S.J. Bultman, C.E. Sims, S.T. Magness, N.L. Allbritton, A microengineered collagen scaffold for generating a polarized crypt-villus architecture of human small intestinal epithelium, Biomaterials 128 (2017) 44–55.

[52] V. De Gregorio, G. Imparato, F. Urciuolo, P.A. Netti, Micro-patterned endogenous stroma equivalent induces polarized crypt-villus architecture of human small intestinal epithelium, Acta Biomater 81 (2018) 43–59.

[53] C.M. Costello, M.B. Phillipsen, L.M. Hartmanis, M.A. Kwasnica, V. Chen, D. Hackam, M.W. Chang, W.E. Bentley, J.C. March, Microscale bioreactors for in situ characterization of gi epithelial cell physiology, Sci Rep 7 (2017) 12515.

[54] Q. Zeng, L. Liu, Y. Guo, R. Li, J. Sun, C. Guo, Preparation of small intestinal submucosa as a scaffold for cardiac tissue engineering, 2011 4th International Conference on Biomedical Engineering and Informatics (BMEI), 2011, pp. 1251–1255.

[55] B.J. Koestler, C.M. Ward, C.R. Fisher, A. Rajan, A.W. Maresso, S.M. Payne, Human intestinal enteroids as a model system of shigella pathogenesis, Infect Immun 87 (2019) e00733–18.

