## Supplementary Tables and Figures for "Effect of Substrate Stiffness on Human Intestinal Enteroids Infectivity by Enteroaggregative *Escherichia coli*"

**Supplemental Table S1.** Statistical p-values from the comparison of the number of adhered bacteria atop J2 HIEs cultured on different substrates (Figure 4B). Kruskal Wallis test (non-parametric data) with Dunn's multiple comparisons.

| | | | |
|---|---|---|---|
| Number of families | 1 | | |
| Number of comparisons per family | 10 | | |
| Alpha | 0.05 | | |
| **Dunn's multiple comparisons test** | **Significant?** | **Summary** | **Adjusted P Value** |
| Glass vs. Plastic | Yes | *** | 0.0002 |
| Glass vs. Stiff | Yes | ** | 0.0066 |
| Glass vs. Medium | Yes | **** | <0.0001 |
| Glass vs. Soft | Yes | **** | <0.0001 |
| Plastic vs. Stiff | No | ns | >0.9999 |
| Plastic vs. Medium | No | ns | >0.9999 |
| Plastic vs. Soft | No | ns | 0.7826 |
| Stiff vs. Medium | No | ns | 0.0984 |
| Stiff vs. Soft | Yes | ** | 0.0016 |
| Medium vs. Soft | No | ns | >0.9999 |

**Supplemental Table S2.** Statistical p-values from the comparison of the mean cluster size of bacterial clusters atop J2 HIEs cultured on different substrates (Figure 4C). Kruskal Wallis test (non-parametric data) with Dunn's multiple comparisons.

| | | | |
|---|---|---|---|
| Number of families | 1 | | |
| Number of comparisons per family | 10 | | |
| Alpha | 0.05 | | |
| **Dunn's multiple comparisons test** | **Significant?** | **Summary** | **Adjusted P Value** |
| Glass vs. Plastic | Yes | * | 0.0242 |
| Glass vs. Stiff | No | ns | 0.4181 |
| Glass vs. Medium | Yes | **** | <0.0001 |
| Glass vs. Soft | Yes | **** | <0.0001 |
| Plastic vs. Stiff | No | ns | >0.9999 |
| Plastic vs. Medium | No | ns | >0.9999 |
| Plastic vs. Soft | No | ns | 0.1869 |
| Stiff vs. Medium | Yes | ** | 0.0079 |
| Stiff vs. Soft | Yes | *** | 0.0001 |
| Medium vs. Soft | No | ns | >0.9999 |

**Supplemental Table S3.** Statistical p-values from the comparison of the number of differently sized bacterial clusters atop J2 HIEs cultured on different substrates (Figure 4D). 2-way ANOVA (parametric data) with Tukey multiple comparisons.

| Within each row, compare columns (simple effects within rows) | | | |
|---|---|---|---|
| Number of families | 7 | | |
| Number of comparisons per family | 10 | | |
| Alpha | 0.05 | | |
| **Tukey's multiple comparisons test** | **Significant?** | **Summary** | **Adjusted P Value** |
| **1 bacteria in cluster** | | | |
| Glass vs. Plastic | No | ns | 0.2813 |
| Glass vs. Stiff | No | ns | 0.1689 |
| Glass vs. Medium | No | ns | 0.1910 |
| Glass vs. Soft | Yes | * | 0.0308 |
| Plastic vs. Stiff | No | ns | >0.9999 |
| Plastic vs. Medium | No | ns | >0.9999 |
| Plastic vs. Soft | No | ns | 0.9837 |
| Stiff vs. Medium | No | ns | >0.9999 |
| Stiff vs. Soft | No | ns | 0.9507 |
| Medium vs. Soft | No | ns | 0.9348 |
| **2 bacteria in cluster** | | | |
| Glass vs. Plastic | No | ns | 0.4936 |
| Glass vs. Stiff | No | ns | 0.4923 |
| Glass vs. Medium | No | ns | 0.3402 |
| Glass vs. Soft | No | ns | 0.0680 |
| Plastic vs. Stiff | No | ns | 0.9992 |
| Plastic vs. Medium | No | ns | >0.9999 |
| Plastic vs. Soft | No | ns | 0.9657 |
| Stiff vs. Medium | No | ns | 0.9990 |
| Stiff vs. Soft | No | ns | 0.8233 |
| Medium vs. Soft | No | ns | 0.9278 |
| **3-10 bacteria in cluster** | | | |
| Glass vs. Plastic | Yes | **** | <0.0001 |
| Glass vs. Stiff | No | ns | 0.1057 |
| Glass vs. Medium | Yes | *** | 0.0003 |
| Glass vs. Soft | Yes | **** | <0.0001 |
| Plastic vs. Stiff | No | ns | 0.0873 |
| Plastic vs. Medium | No | ns | 0.8570 |
| Plastic vs. Soft | No | ns | 0.9572 |
| Stiff vs. Medium | No | ns | 0.3659 |
| Stiff vs. Soft | Yes | ** | 0.0015 |
| Medium vs. Soft | No | ns | 0.2680 |

| | | | |
|---|---|---|---|
| **11-50 bacteria in cluster** | | | |
| Glass vs. Plastic | No | ns | 0.2691 |
| Glass vs. Stiff | No | ns | 0.8020 |
| Glass vs. Medium | No | ns | 0.1204 |
| Glass vs. Soft | Yes | **** | <0.0001 |
| Plastic vs. Stiff | No | ns | 0.8152 |
| Plastic vs. Medium | No | ns | >0.9999 |
| Plastic vs. Soft | No | ns | 0.2752 |
| Stiff vs. Medium | No | ns | 0.6744 |
| Stiff vs. Soft | Yes | ** | 0.0029 |
| Medium vs. Soft | No | ns | 0.1413 |
| **51-100 bacteria in cluster** | | | |
| Glass vs. Plastic | No | ns | >0.9999 |
| Glass vs. Stiff | No | ns | >0.9999 |
| Glass vs. Medium | No | ns | 0.4357 |
| Glass vs. Soft | No | ns | 0.0983 |
| Plastic vs. Stiff | No | ns | >0.9999 |
| Plastic vs. Medium | No | ns | 0.5745 |
| Plastic vs. Soft | No | ns | 0.1952 |
| Stiff vs. Medium | No | ns | 0.4798 |
| Stiff vs. Soft | No | ns | 0.1083 |
| Medium vs. Soft | No | ns | 0.9237 |
| **>100 bacteria in cluster** | | | |
| Glass vs. Plastic | No | ns | >0.9999 |
| Glass vs. Stiff | No | ns | >0.9999 |
| Glass vs. Medium | Yes | ** | 0.0067 |
| Glass vs. Soft | Yes | **** | <0.0001 |
| Plastic vs. Stiff | No | ns | >0.9999 |
| Plastic vs. Medium | Yes | * | 0.0268 |
| Plastic vs. Soft | Yes | **** | <0.0001 |
| Stiff vs. Medium | Yes | ** | 0.0042 |
| Stiff vs. Soft | Yes | **** | <0.0001 |
| Medium vs. Soft | Yes | **** | <0.0001 |

**Supplemental Table S4.** Statistically signficant p-values from the comparison of the number of adhered bacteria and bacterial cluster size atop J11 HIEs cultured on different substrates (Supplemental Figures S2A-C). Statistical analyses performed as in previous tables.

| **Analysis and significant comparison** | **Adjusted p-value** |
|---|---|
| Number of adherent bacteria per image | |
| Stiff vs. Glass | 0.0238 |
| Mean cluster size per image | |
| Soft vs. Glass | 0.0439 |
| Stiff vs. Glass | 0.0287 |
| Size of every cluster across all images | |
| Soft vs. Glass | 0.0001 |
| Stiff vs. Glass | 0.0004 |

**Supplemental Table S5.** Statistical p-values from the comparison of the number of differently sized bacterial clusters atop J11 HIEs cultured on different substrates (Supplemental Figure 2D). 2-way ANOVA (parametric data) with Tukey multiple comparisons.

| Within each row, compare columns (simple effects within rows) | | | |
|---|---|---|---|
| Number of families | 6 | | |
| Number of comparisons per family | 6 | | |
| Alpha | 0.05 | | |
| **Tukey's multiple comparisons test** | **Significant?** | **Summary** | **Adjusted P Value** |
| **1 bacteria in cluster** | | | |
| Glass vs. Stiff | No | ns | 0.6193 |
| Glass vs. Medium | No | ns | >0.9999 |
| Glass vs. Soft | No | ns | 0.5989 |
| Stiff vs. Medium | No | ns | 0.7497 |
| Stiff vs. Soft | No | ns | >0.9999 |
| Medium vs. Soft | No | ns | 0.7350 |
| **2 bacteria in cluster** | | | |
| Glass vs. Stiff | No | ns | 0.6040 |
| Glass vs. Medium | No | ns | 0.9498 |
| Glass vs. Soft | No | ns | 0.9407 |
| Stiff vs. Medium | No | ns | 0.4350 |
| Stiff vs. Soft | No | ns | 0.9094 |
| Medium vs. Soft | No | ns | 0.7583 |
| **3-10 bacteria in cluster** | | | |
| Glass vs. Stiff | Yes | **** | <0.0001 |
| Glass vs. Medium | No | ns | 0.2669 |
| Glass vs. Soft | Yes | ** | 0.0059 |
| Stiff vs. Medium | No | ns | 0.1488 |
| Stiff vs. Soft | No | ns | 0.3231 |

| | | | |
|---|---|---|---|
| Medium vs. Soft | No | ns | 0.8599 |
| **11-50 bacteria in cluster** | | | |
| Glass vs. Stiff | Yes | * | 0.0175 |
| Glass vs. Medium | No | ns | >0.9999 |
| Glass vs. Soft | Yes | ** | 0.0026 |
| Stiff vs. Medium | No | ns | 0.1003 |
| Stiff vs. Soft | No | ns | 0.9324 |
| Medium vs. Soft | Yes | * | 0.0308 |
| **51-100 bacteria in cluster** | | | |
| Glass vs. Stiff | No | ns | >0.9999 |
| Glass vs. Medium | No | ns | 0.9909 |
| Glass vs. Soft | No | ns | 0.8989 |
| Stiff vs. Medium | No | ns | 0.9925 |
| Stiff vs. Soft | No | ns | 0.9079 |
| Medium vs. Soft | No | ns | 0.9946 |
| **100-1000 bacteria in cluster** | | | |
| Glass vs. Stiff | No | ns | 0.5149 |
| Glass vs. Medium | No | ns | >0.9999 |
| Glass vs. Soft | No | ns | 0.9621 |
| Stiff vs. Medium | No | ns | 0.7016 |
| Stiff vs. Soft | No | ns | 0.8091 |
| Medium vs. Soft | No | ns | 0.9811 |

**Supplemental Table S6.** Statistically signficant p-values from the comparison of the number of adhered bacteria and bacterial cluster size atop D109 HIEs cultured on different substrates (Supplemental Figures S3A and C). Statistical analyses performed as in previous tables.

| **Analysis and significant comparison** | **Adjusted p-value** |
|---|---|
| Number of adherent bacteria per image | |
| Stiff vs. Soft | 0.0077 |
| Size of every cluster across all images | |
| Glass vs. Soft | <0.0001 |
| Medium vs. Soft | <0.0001 |

**Supplemental Table S7.** Statistical p-values from the comparison of the number of differently sized bacterial clusters atop D109 HIEs cultured on different substrates (Supplemental Figure 3D). 2-way ANOVA (parametric data) with Tukey multiple comparisons.

| | | | |
|---|---|---|---|
| Within each row, compare columns (simple effects within rows) | | | |
| Number of families | 7 | | |
| Number of comparisons per family | 6 | | |
| Alpha | 0.05 | | |
| **Tukey's multiple comparisons test** | **Significant?** | **Summary** | **Adjusted P Value** |
| **1 bacteria in cluster** | | | |
| Glass vs. Stiff | No | ns | 0.9942 |
| Glass vs. Medium | No | ns | >0.9999 |
| Glass vs. Soft | No | ns | 0.9799 |
| Stiff vs. Medium | No | ns | 0.9946 |
| Stiff vs. Soft | No | ns | 0.9440 |
| Medium vs. Soft | No | ns | 0.9856 |
| **2 bacteria in cluster** | | | |
| Glass vs. Stiff | No | ns | 0.9908 |
| Glass vs. Medium | No | ns | 0.9996 |
| Glass vs. Soft | No | ns | 0.9998 |
| Stiff vs. Medium | No | ns | 0.9981 |
| Stiff vs. Soft | No | ns | 0.9986 |
| Medium vs. Soft | No | ns | >0.9999 |
| **3-10 bacteria in cluster** | | | |
| Glass vs. Stiff | No | ns | 0.9551 |
| Glass vs. Medium | No | ns | 0.9999 |
| Glass vs. Soft | No | ns | 0.9994 |
| Stiff vs. Medium | No | ns | 0.9772 |
| Stiff vs. Soft | No | ns | 0.9918 |
| Medium vs. Soft | No | ns | >0.9999 |
| **11-50 bacteria in cluster** | | | |
| Glass vs. Stiff | No | ns | 0.9829 |
| Glass vs. Medium | No | ns | 0.9999 |
| Glass vs. Soft | No | ns | 0.9987 |
| Stiff vs. Medium | No | ns | 0.9934 |
| Stiff vs. Soft | No | ns | 0.9990 |
| Medium vs. Soft | No | ns | 0.9998 |
| **51-100 bacteria in cluster** | | | |
| Glass vs. Stiff | No | ns | 0.9990 |
| Glass vs. Medium | No | ns | 0.9975 |
| Glass vs. Soft | No | ns | 0.9873 |
| Stiff vs. Medium | No | ns | 0.9908 |

| | | | |
|---|---|---|---|
| Stiff vs. Soft | No | ns | 0.9756 |
| Medium vs. Soft | No | ns | 0.9985 |
| **100-1000 bacteria in cluster** | | | |
| Glass vs. Stiff | No | ns | 0.9875 |
| Glass vs. Medium | No | ns | 0.2591 |
| Glass vs. Soft | No | ns | 0.1858 |
| Stiff vs. Medium | No | ns | 0.2068 |
| Stiff vs. Soft | No | ns | 0.1466 |
| Medium vs. Soft | No | ns | 0.9674 |
| **>1000 bacteria in cluster** | | | |
| Glass vs. Stiff | No | ns | >0.9999 |
| Glass vs. Medium | No | ns | >0.9999 |
| Glass vs. Soft | Yes | **** | <0.0001 |
| Stiff vs. Medium | No | ns | >0.9999 |
| Stiff vs. Soft | Yes | **** | <0.0001 |
| Medium vs. Soft | Yes | **** | <0.0001 |

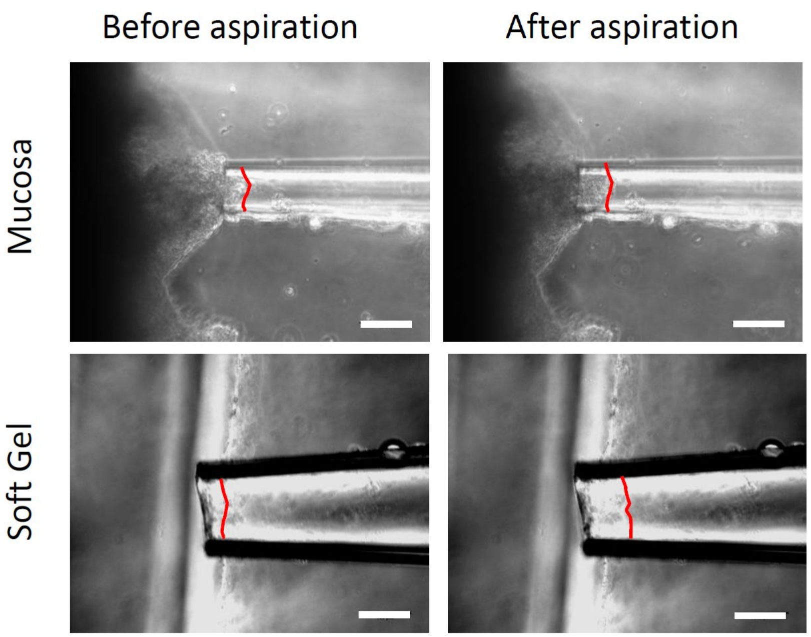

**Supplemental Fig. S1.** Images of micropipette aspiration of the intestinal mucosa and soft hydrogel. Red line indicates upper boundary of aspirated material within the micropipette. White scale bar = 100 microns.

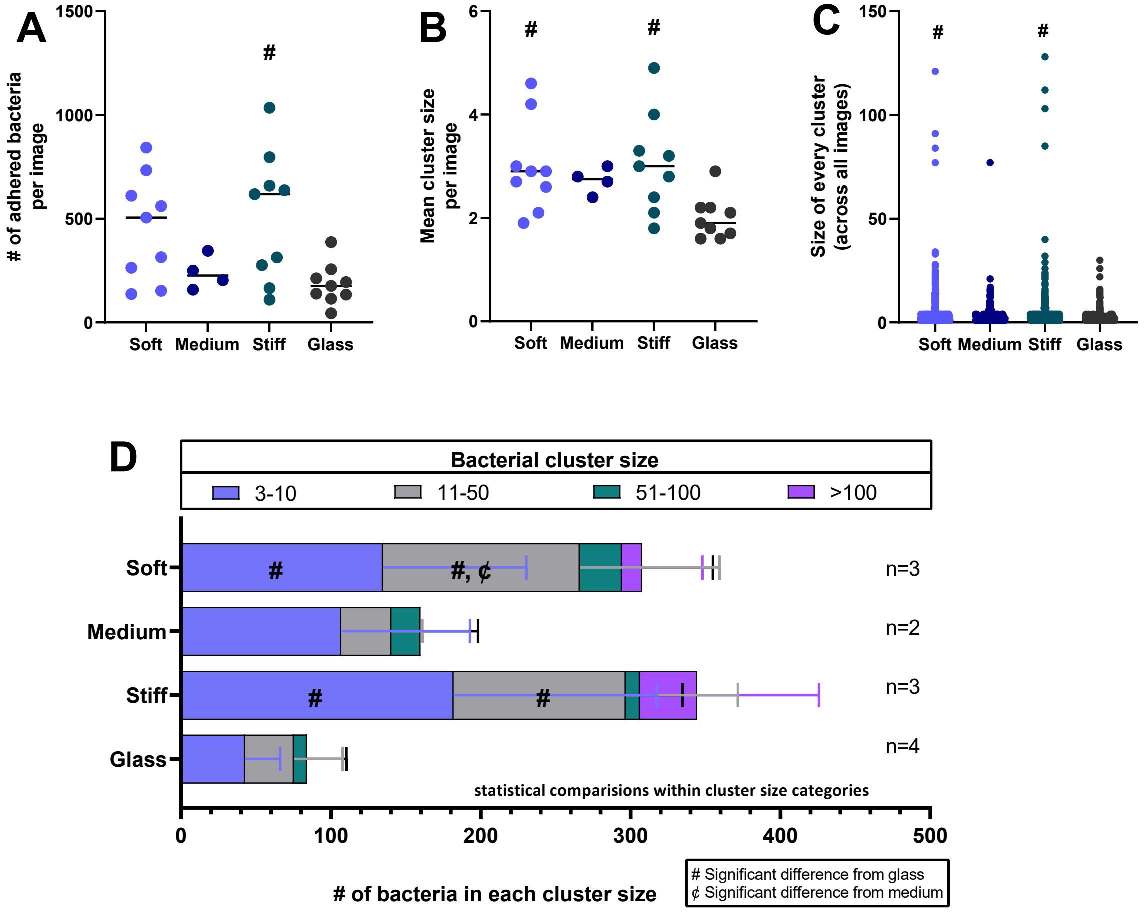

**Supplemental Fig. S2.** *Adherence and aggregation pattern of EAEC 042 on J11 monolayers plated on hydrogels of varying stiffness.* J11 monolayers were plated on hydrogels or on glass chambered slide controls, differentiated 5 days, and infected for 3 hours with 042 at an MOI=10. Samples were fixed and stained using a Hema3 kit. For each well, 3 images were taken and counted (e.g., n=3 has data from a total of 9 images). All comparisons were made with Tukey's multiple comparisons test following one-way ANOVA. For simplicity, clusters of 1-2 bacteria are not shown (numbers of clusters of 1-2 bacteria were not significantly different between groups), but were included in the statistical analysis shown in Supplemental Table S5.

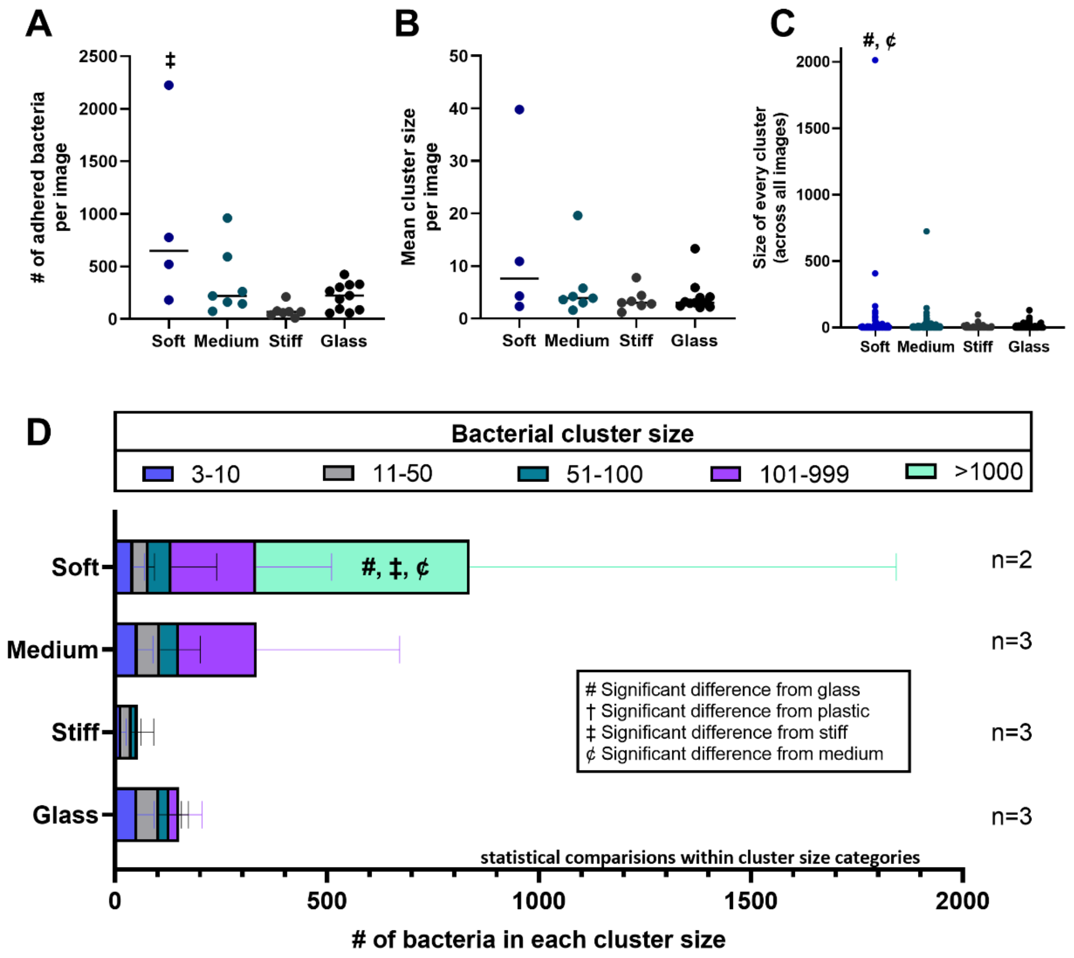

**Supplemental Fig. S3.** *Adherence and aggregation pattern of EAEC 042 on D109 monolayers plated on hydrogels of varying stiffness.* D109 monolayers were plated on hydrogels or on glass chambered slide controls, differentiated 5 days, and infected for 3 hours with 042 at an MOI=10. Samples were fixed and stained using a Hema3 kit. For each well, 3 images were taken and counted (e.g., n=3 has data from a total of 9 images). All comparisons were made with Tukey's multiple comparisons test following one-way ANOVA. For simplicity, clusters of 1-2 bacteria are not shown (numbers of clusters of 1-2 bacteria were not significantly different between groups), but were included in the statistical analysis shown in Supplemental Table S7.

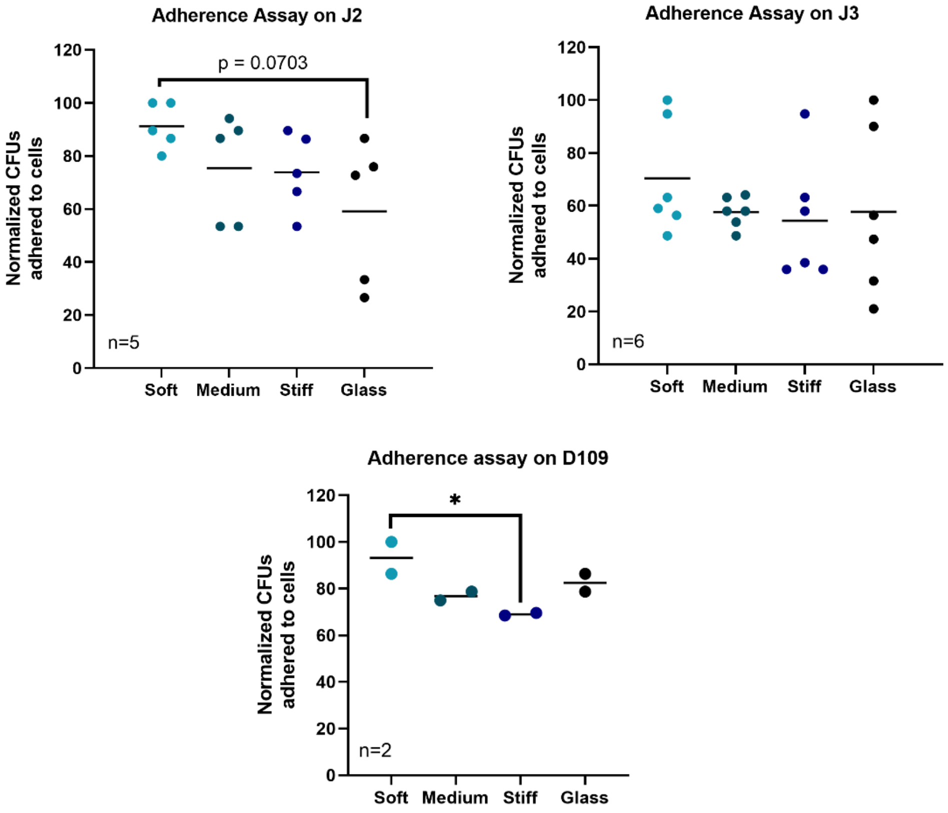

**Supplemental Fig. S4.** *Adherence of Enteroaggregative E. coli to jejunal and duodenal monolayers grown on hydrogels of various stiffnesses.* J2, J3, and D109 HIE monolayers were cultured with prototype EAEC strain 042 for 3 hours, washed to remove unadhered bacteria, and scraped off the plate. The cell solution was then serially diluted, plated on LB plates, and counted for CFUs. Data shown was calculated from one CFU for each well. n=2-6 wells per condition as indicated on the chart. * denotes $p \leq 0.05$ between groups.

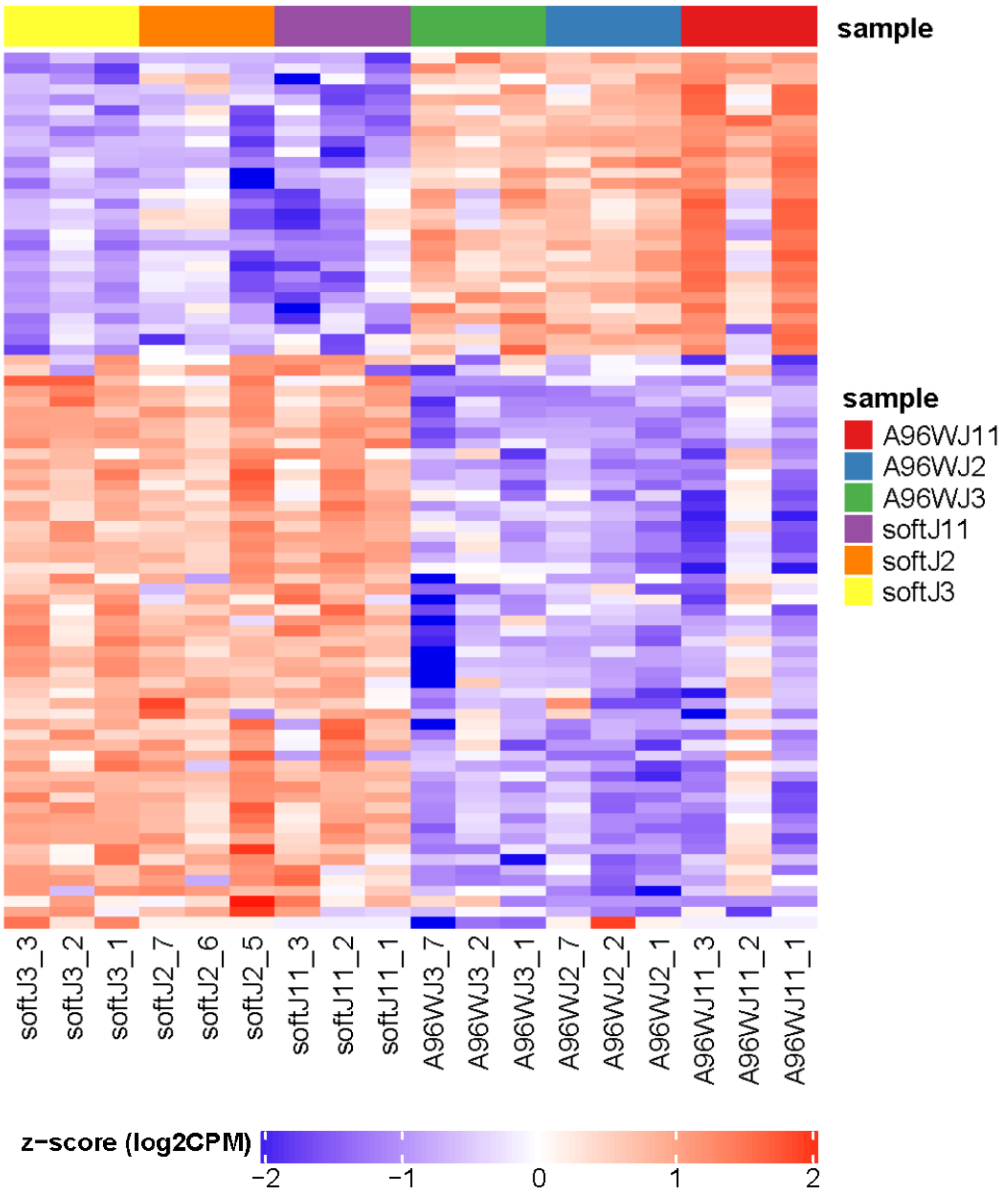

**Supplemental Fig. S5.** Heat map for selected genes from biological pathways, known to be mechanosensitive, that were identified through Gene Set Enrichment Analysis. All genes shown have an FDR < 0.05 and |log FC| > 2. Left columns show gene expression in the soft hydrogels and right columns show the 96-well plates. Red = upregulated, blue = downregulated. The heat maps for the medium and stiff hydrogels vs. 96-well plates were very similar. A subset of this heat map is shown in Fig. 6D. Data and gene loci for this heatmap and the smaller version are found in Supplemental Data Set 2.

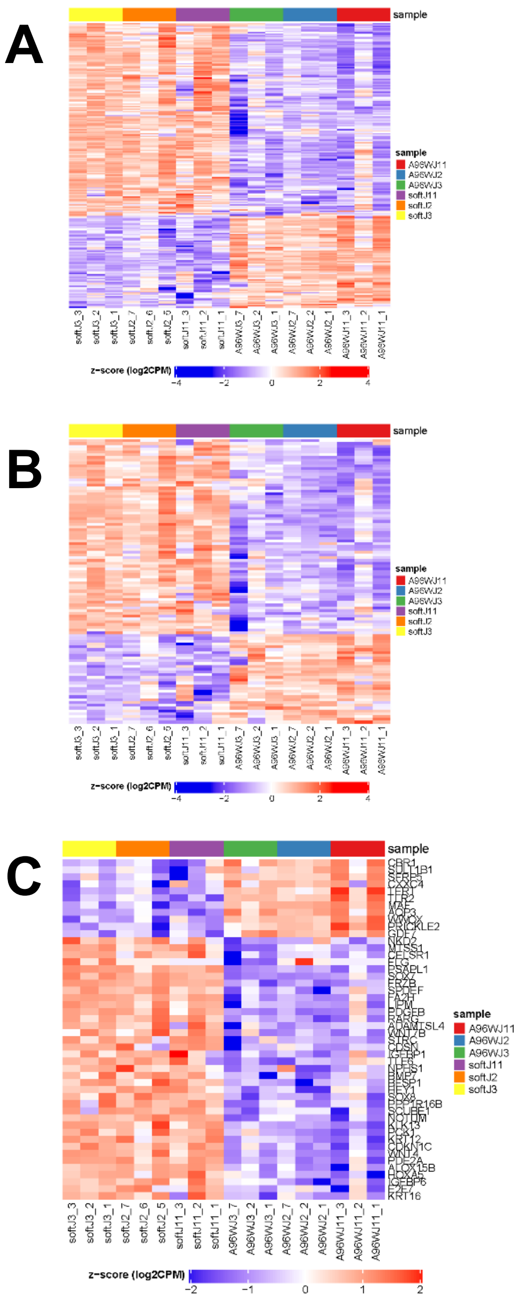

**Supplemental Fig. S6.** Heat maps relevant to epithelial differentiation and Wnt signaling. All genes shown have an FDR < 0.05. Left columns show gene expression in the soft hydrogels and right columns show the 96-well plates. Red = upregulated, blue = downregulated. The heat maps for the medium and stiff hydrogels vs. 96-well plates were very similar. Data and gene loci for the heatmaps are found in Supplemental Data Set 2. **(A)** Heat map of Gene Ontology Epithelial Cell Differentiation pathway. **(B)** Heat map of Gene Ontology Canonical Wnt Signaling pathway. **(C)** Consolidated heat map for genes from multiple epithelial differentiation and Wnt pathways with high Normalized Enrichment Scores that were identified through Gene Set Enrichment Analysis. All genes shown in this consolidated heat map have |log FC| > 2.0. A subset of this heat map is shown in Fig. 7A.

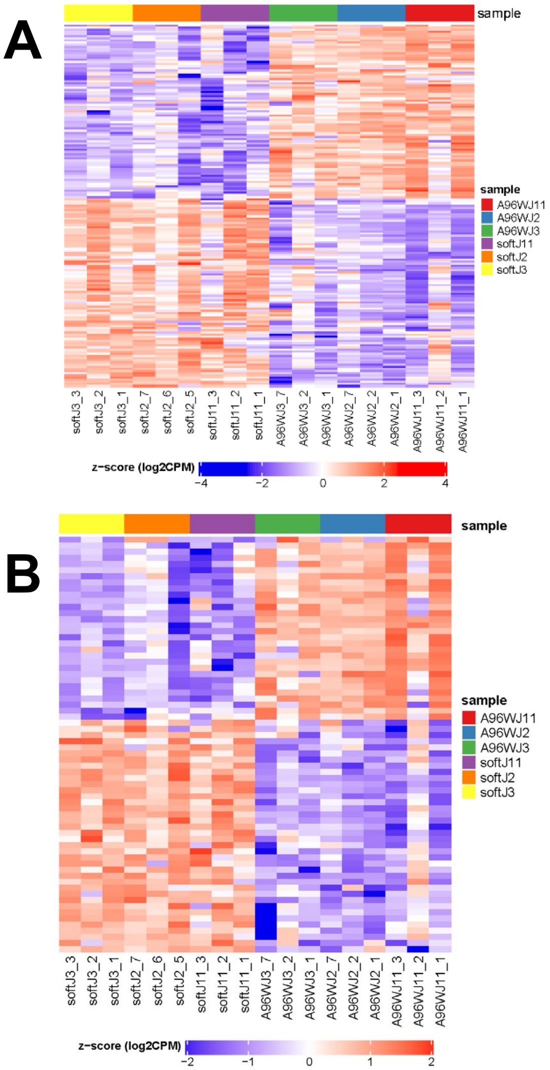

**Supplemental Fig. S7.** Heat maps relevant to bacteria and proteoglycans. All genes shown have an FDR < 0.05. Left columns show gene expression in the soft hydrogels and right columns show the 96-well plates. Red = upregulated, blue = downregulated. The heat maps for the medium and stiff hydrogels vs. 96-well plates were very similar. Data and gene loci for the heatmaps are found in Supplemental Data Set 2. **(A)** Heat map of Gene Ontology Response to Bacterium pathway. **(B)** Consolidated heat map for genes from multiple bacterial-related and proteoglycan-related pathways with high Normalized Enrichment Scores that were identified through Gene Set Enrichment Analysis. All genes shown in this consolidated heat map have |log FC| > 2.0. A subset of this heat map is shown in Fig. 7B.

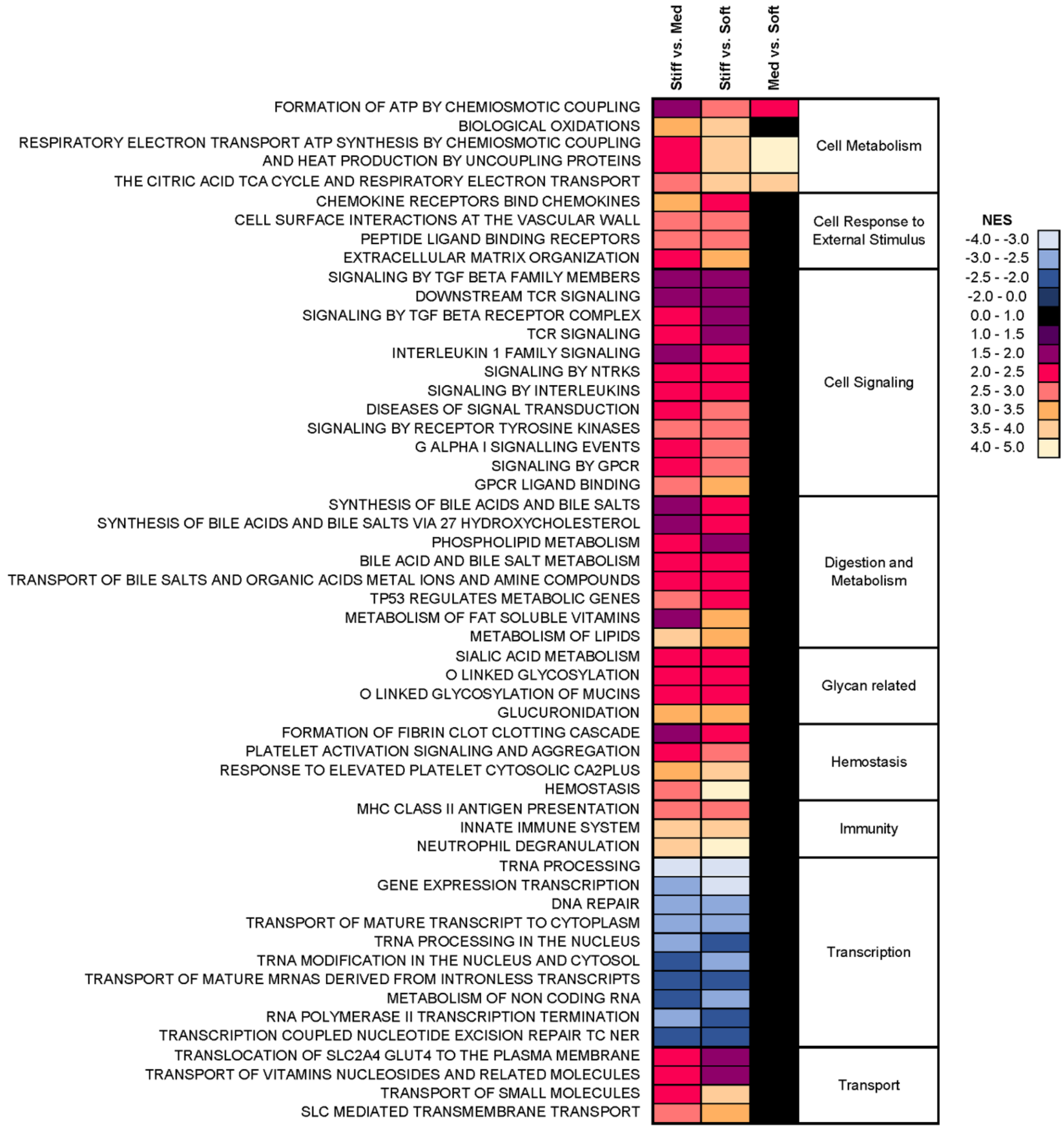

**Supplemental Fig. S8.** Normalized enrichment scores (NES) of selected Reactome pathways that were significantly different (FDR < 0.05) between HIEs cultured atop hydrogels of different stiffness (Soft, Medium, Stiff). In each column, warmer colors indicate greater enrichment (positive correlation with 1st gel stiffness shown) and cooler colors indicate least enrichment (positive correlation with 2nd gel stiffness shown). Data were averaged across the 3 patients.

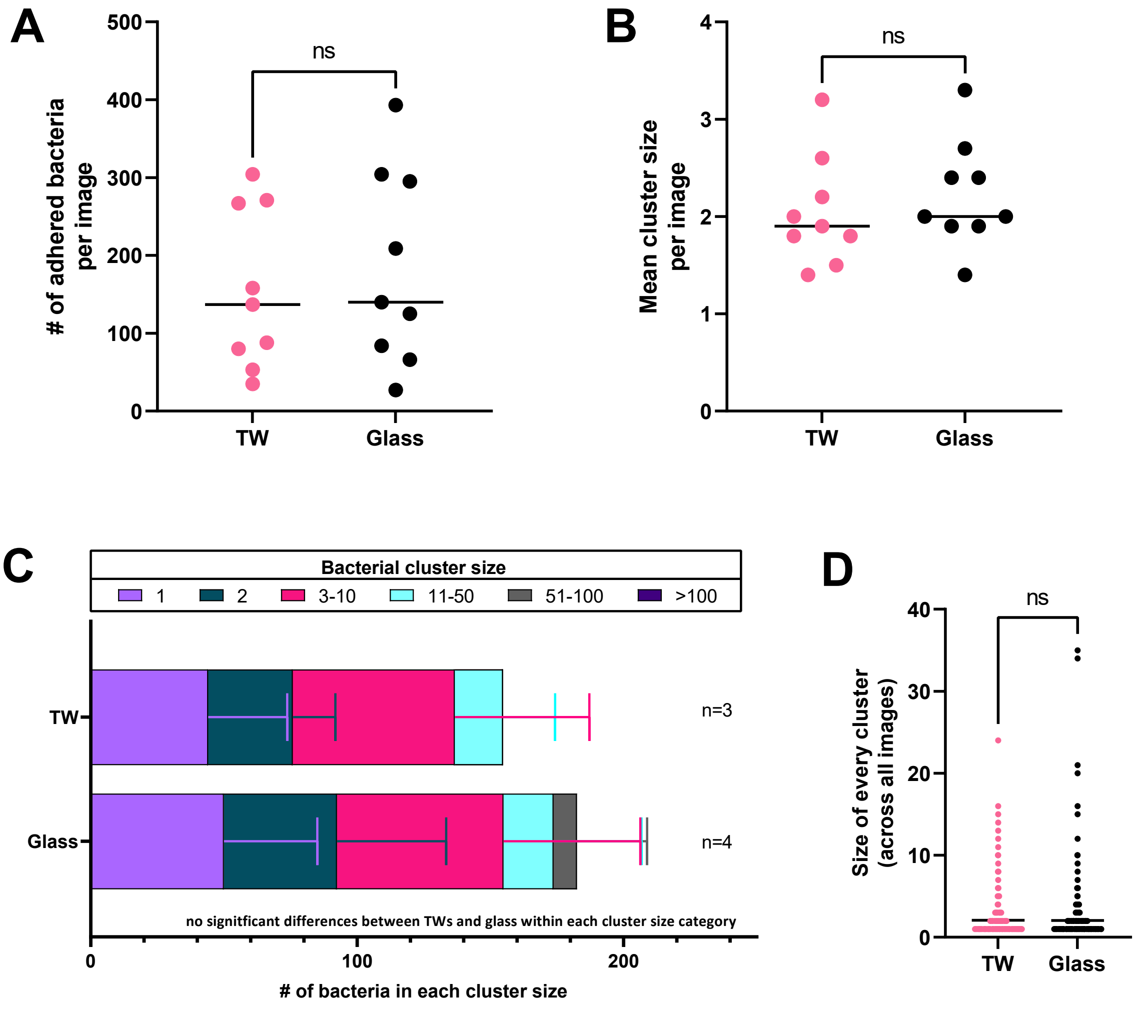

**Supplemental Fig. S9.** *Adherence and aggregation pattern of EAEC 042 on J2 monolayers plated on Transwells (TW) and glass chambered slides.* J2 monolayers were differentiated 5 days previous to 2 hour infection with 042 at an MOI=10. Samples were fixed and stained using a Hema3 kit. For each well, 3 images were taken and counted (e.g. n=3 has data from a total of 9 images). Enteroids plated on Transwells were compared to those plated on glass using an unpaired t-test.

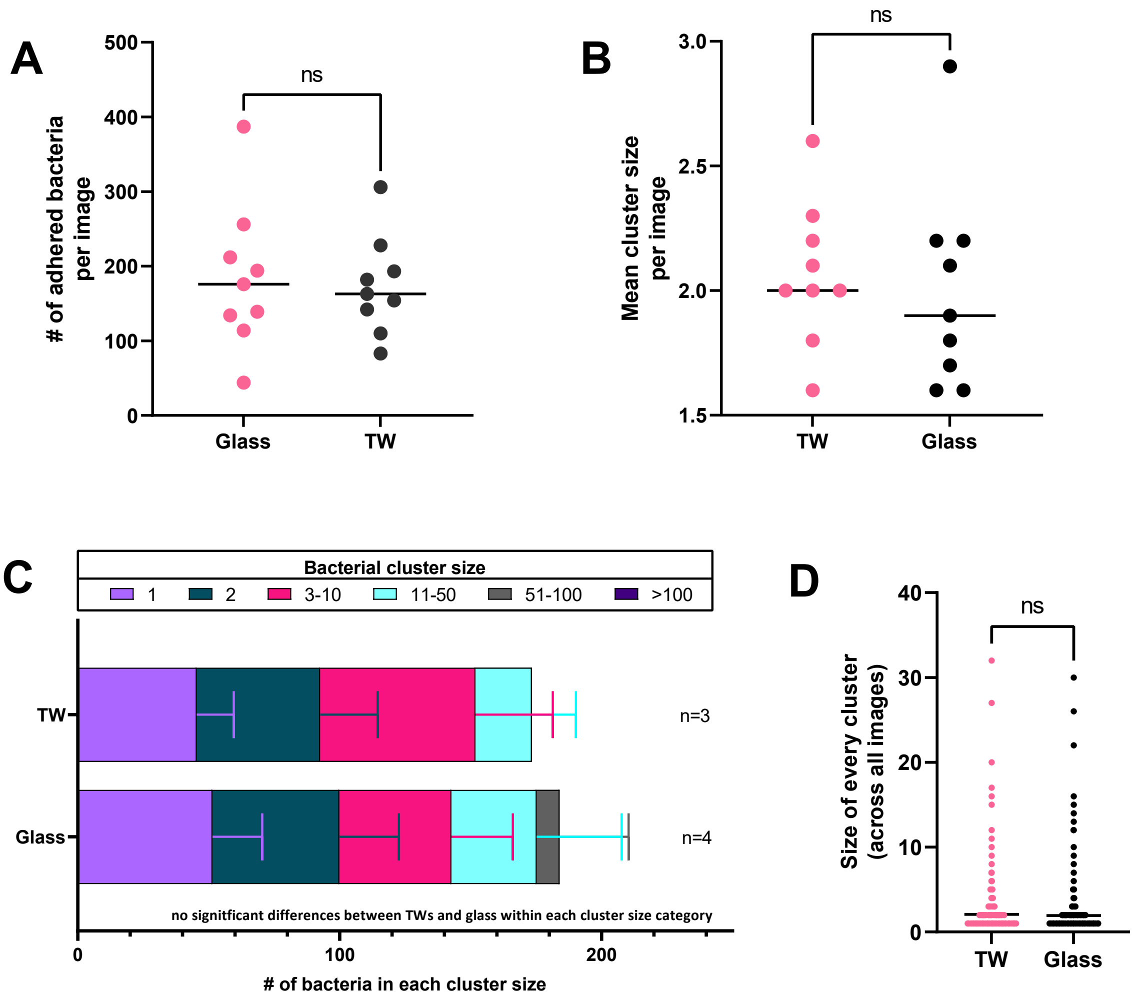

**Supplemental Fig. 10.** *Adherence and aggregation pattern of EAEC 042 on J11 monolayers plated on transwells and glass chambered slides.* J11 monolayers were differentiated 5 days previous to 2 hour infection with 042 at an MOI=10. Samples were fixed and stained using a Hema3 kit. For each well, 3 images were taken and counted (e.g., n=3 has data from a total of 9 images). Enteroids plated on Transwells were compared to those plated on glass using an unpaired t-test.

**Supplemental Data Set 1.** Set of up-regulated and down-regulated genes in HIEs.
The excel sheets list all the different genes that are up- or down-regulated in HIEs grown atop soft, medium, or stiff hydrogels vs. HIEs grown atop 96-well plates. Data are averaged from 3 patients. Additional sheets provide the names of genes (with log FC > 2.0) organized as in the Venn diagrams.

**Supplemental Data Set 2.** Gene data from heat maps.
The excel sheets are numerical versions of the heat maps shown in Figures 6-7 and Supplemental Figures S1-S3.

**Supplemental Data Set 3.** List of all GSEA pathways.

The excel sheets list normalized enrichment scores (NES) for significantly enriched biological pathways. Data are averaged from three patients. All pathways listed had a nonzero NES for at least one of the six comparisons (hydrogel vs. 96-well plate or between-hydrogels).
